# Extracellular matrix-inducing Sox9 orchestrates basal progenitor proliferation and gliogenesis in developing neocortex

**DOI:** 10.1101/704890

**Authors:** Ayse Güven, Denise Stenzel, Katherine R. Long, Marta Florio, Holger Brandl, Wieland B. Huttner

## Abstract

Neocortex expansion is largely based on the proliferative capacity of basal progenitors (BPs), which is increased by extracellular matrix (ECM) components via integrin signaling. Here we show that Sox9 drives expression of ECM components and that laminin 211 increases BP proliferation in embryonic mouse neocortex. Examination of Sox9 expression reveals that Sox9 is expressed in BPs of developing ferret and human, but not mouse neocortex. Functional studies by conditional Sox9 expression in the mouse BP lineage demonstrate increased BP proliferation, reduced Tbr2 and induction of Olig2 expression, indicative of premature gliogenesis. Conditional Sox9 expression also results in cell non-autonomous stimulation of BP proliferation followed by increased production of upper-layer neurons. Collectively, our findings demonstrate that Sox9 exerts concerted effects on transcription, BP proliferation, neuron production, and neurogenic as well as gliogenic BP cell fate, suggesting that Sox9 acts a master regulator in the subventricular zone to promote neocortical expansion.

## Introduction

The expansion of the neocortex in the course of human evolution and its growth during the development of the human brain have become a central topic of molecularly and cellularly focused developmental neuroscience. Neocortical expansion is thought to constitute one of the bases for the unique cognitive abilities of humans. At the cellular level, the proliferative capacity and pool size of cortical neural progenitor cells (cNPCs) is regarded as a key parameter underlying neocortical expansion (Arlotta and Pasca, 2019; Borrell and Reillo, 2012; Dehay et al., 2015; Fernandez et al., 2016; Fietz and Huttner, 2011; Fish et al., 2008; Florio and Huttner, 2014; Kriegstein et al., 2006; Lui et al., 2011; Martynoga et al., 2012; Miller et al., 2019; Mitchell and Silver, 2018; Namba and Huttner, 2017; Rakic, 2009; Silbereis et al., 2016; Uzquiano et al., 2018).

There are two major germinal zones in the developing neocortex, which harbor two principal classes of cNPCs that exhibit distinct features related to the apical-basal polarity of the cortical wall. The apical-most, and primary, germinal zone, the ventricular zone (VZ), harbors the cell bodies of cNPCs collectively referred to as apical progenitors, all of which contact the ventricle (Taverna et al., 2014). Among these, apical (or ventricular) radial glia (aRG), which arise by transition from neuroepithelial cells as the generation of neocortical neurons begins (Götz and Huttner, 2005; Kriegstein and Götz, 2003) and which exhibit pronounced apical-basal cell polarity (Götz and Huttner, 2005; Reiner et al., 2012; Taverna et al., 2014), have been recognized as a major cNPC type (Lui et al., 2011; Namba and Huttner, 2017). The zone basal to the VZ, the subventricular zone (SVZ), constitutes a secondary germinal layer that harbors the cell bodies of cNPCs collectively referred to as basal progenitors (BPs), all of which lack contact with the ventricle (Borrell and Reillo, 2012; Dehay et al., 2015; Fietz and Huttner, 2011; Kriegstein and Alvarez-Buylla, 2009; Pontious et al., 2008; Taverna et al., 2014). There are two main types of BPs, (i) basal intermediate progenitors (bIPs), which lack apical-basal cell polarity and do not exhibit significant cell processes at mitosis (Attardo et al., 2008; Haubensak et al., 2004; Miyata et al., 2004; Noctor et al., 2004), and (ii) basal (or outer) radial glia (bRG), which exhibit basal and/or apical cell polarity, extending basal and/or apically directed cell processes throughout their cell cycle including mitosis (Betizeau et al., 2013; Fietz et al., 2010; Hansen et al., 2010; Pilz et al., 2013; Reillo et al., 2011; Shitamukai et al., 2011; Wang et al., 2011).

For cell biological reasons related to their apical cell polarity, aRG mitoses are confined to the ventricular surface, a limited space, which poses a constraint with regard to maximizing their number and, consequently, to increasing aRG pool size (Fietz and Huttner, 2011; Fish et al., 2008; Taverna et al., 2014). In contrast, this constraint does not exist for BPs. By virtue of these cells having delaminated from the ventricular surface, BPs have an intrinsic advantage compared to aRG with regard to maximizing the number of their mitoses and hence to increasing their pool size, as they can undergo mitosis virtually anywhere along the radial axis of the SVZ (Betizeau et al., 2013; Fietz and Huttner, 2011; Fietz et al., 2010; Fish et al., 2008; Florio and Huttner, 2014; Hansen et al., 2010; Reillo et al., 2011). Accordingly, neocortical expansion is thought to be linked to an increase in the proliferative capacity of BPs, resulting in their increased pool size and a thickening of the SVZ (Borrell and Reillo, 2012; Dehay et al., 2015; Fernandez et al., 2016; Fietz and Huttner, 2011; Florio and Huttner, 2014; Kriegstein et al., 2006; Lui et al., 2011). Thus, in mammals lacking neocortical expansion, such as mouse, BPs exhibit only low proliferative capacity, typically dividing only once to generate two post-mitotic neurons, and their pool size is relatively small and the SVZ comparably thin (Arai et al., 2011; Florio and Huttner, 2014; Haubensak et al., 2004; Miyata et al., 2004; Noctor et al., 2004). In contrast, in mammals showing neocortical expansion, notably human, BPs exhibit high proliferative capacity, resulting in an increased pool size and a thick SVZ (Betizeau et al., 2013; Fernandez et al., 2016; Fietz et al., 2010; Florio and Huttner, 2014; Hansen et al., 2010; Pilz et al., 2013; Reillo et al., 2011).

Hence, the crucial question is: what underlies the differences in BP proliferative capacity across the various mammals? An important clue pointing to the differential expression of extracellular matrix (ECM) components as a key regulatory parameter has come from comparative analyses of the transcriptomes of mouse vs. human VZ and SVZ as well as specific cNPC subpopulations. Specifically, in embryonic mouse neocortex, BPs down-regulate the endogenous expression of ECM components in comparison with aRG, in line with the lower proliferative capacity of the former than the latter (Arai et al., 2011). Consistent with this, the expression of ECM components is down-regulated in the embryonic mouse SVZ compared to the VZ, whereas in fetal human neocortex this expression is maintained not only in the VZ, but also inner SVZ (ISVZ) and outer SVZ (OSVZ) (Fietz et al., 2012). Accordingly, not only human aRG, but also human bRG show a characteristic expression of ECM components (Florio et al., 2015; Pollen et al., 2015). Moreover, when mimicking the physiological situation in the fetal human SVZ in an embryonic mouse model, the targeted activation of integrins, the canonical receptors for ECM components, specifically of integrin *α*v*β*3, on mouse BPs promotes their proliferation (Stenzel et al., 2014). Conversely, inhibition of integrin *α*v*β*3 in an *ex vivo* model of the developing neocortex of the ferret, which exhibits an expanded, folded cerebral cortex, reduces the BP pool size, in particular that of bRG (Fietz et al., 2010). Taken together, these findings have led to the concept that an increased expression of ECM components by BPs promotes, via increased integrin signaling, their proliferation (Fietz et al., 2010; Fietz et al., 2012; Kalebic et al., 2019; Long and Huttner, 2019; Stenzel et al., 2014).

This in turn leads to the key question: which transcriptional machinery governs the differential expression of ECM components in the neocortical SVZ of the various mammals? An *in silico* analysis of transcription factors expressed in embryonic mouse vs. fetal human germinal zones and predicted to bind to promoters of ECM genes has revealed Sox9 as a promising candidate to drive expression of ECM components in the human SVZ (Fietz et al., 2012). Here, we have studied the physiological expression of Sox9 in the VZ vs. SVZ of embryonic mouse, embryonic ferret and fetal human neocortex and, based on the results obtained, examined the effects of conditional Sox9 expression in mouse BPs on their proliferative capacity and on driving the expression of ECM genes.

In this context, given the role of Sox9 in driving glia-specific gene expression (Huang et al., 2015; Kang et al., 2012; Klum et al., 2018; Martini et al., 2013; Molofsky et al., 2013; Nagao et al., 2016; Selvaraj et al., 2017) and in the neurogenesis-to-gliogenesis switch in the developing spinal cord (Finzsch et al., 2008; Molofsky et al., 2013; Stolt et al., 2003; Stolt and Wegner, 2010; Wegner and Stolt, 2005), a related issue concerns the neuronal vs. glial fate diversity of neocortical SVZ progenitors across species. In the developing mouse neocortex, neurogenesis and gliogenesis take place sequentially, that is, after neurogenesis, SVZ progenitors of oligodendrocytes and astrocytes are being generated (Kriegstein and Alvarez-Buylla, 2009; Merkle et al., 2004). In contrast, in the SVZ of gyrencephalic species, such as human, macaque and ferret, neurogenic and gliogenic progenitors co-exist already at the peak of neurogenesis (Martinez-Cerdeno et al., 2012; Rash et al., 2019; Reillo and Borrell, 2012; Reillo et al., 2011; Zecevic et al., 2005). We therefore have also examined the effects of conditional Sox9 expression in mouse BPs on their neurogenic vs. gliogenic fate. Taken together, our data provide novel insight into the cell-autonomous vs. cell non-autonomous stimulation of BP proliferation, the consequences for neuron production, and the relationship between neurogenesis and gliogenesis.

## Results

### Sox9-expressing BPs occur in the SVZ of embryonic ferret and fetal human, but not embryonic mouse, neocortex

To gain an initial insight into Sox9 expression in developing neocortex, we analyzed the FPKM values for *Sox9/SOX9* mRNA in the germinal zones and in specific cNPC types of mouse and human neocortex at mid-neurogenesis, using two published transcriptome datasets (Fietz et al., 2012; Florio et al., 2015). In the germinal zones of embryonic day (E) 14.5 mouse neocortex, *Sox9* mRNA was found to be expressed in the VZ, but not SVZ (Supp Fig. 1A, left), whereas in the 13-16 weeks post conception (wpc) human neocortex, *SOX9* mRNA was found to be expressed not only in the VZ, but also in the ISVZ and OSVZ (Supp Fig. 1A, right) (Fietz et al., 2012). In both species, no significant *Sox9/SOX9* mRNA expression was observed in the cortical plate (CP). In specific cNPC types isolated from E14.5 mouse neocortex, *Sox9* mRNA was highly expressed in aRG, but not in bRG, bIPs and neurons (Supp Fig. 1B, left) (Florio et al., 2015). Within mouse aRG, *Sox9* mRNA levels were almost three times as high in the proliferative aRG subpopulation that lacks *Tis21*-GFP expression than in the neurogenic aRG subpopulation that exhibits *Tis21*-GFP expression. In contrast to mouse, in specific cNPC types isolated from 13 wpc human neocortex, *SOX9* mRNA was highly expressed in aRG, but was also found in bRG (Supp Fig. 1B, right) (Florio et al., 2015).

**Fig. 1.**
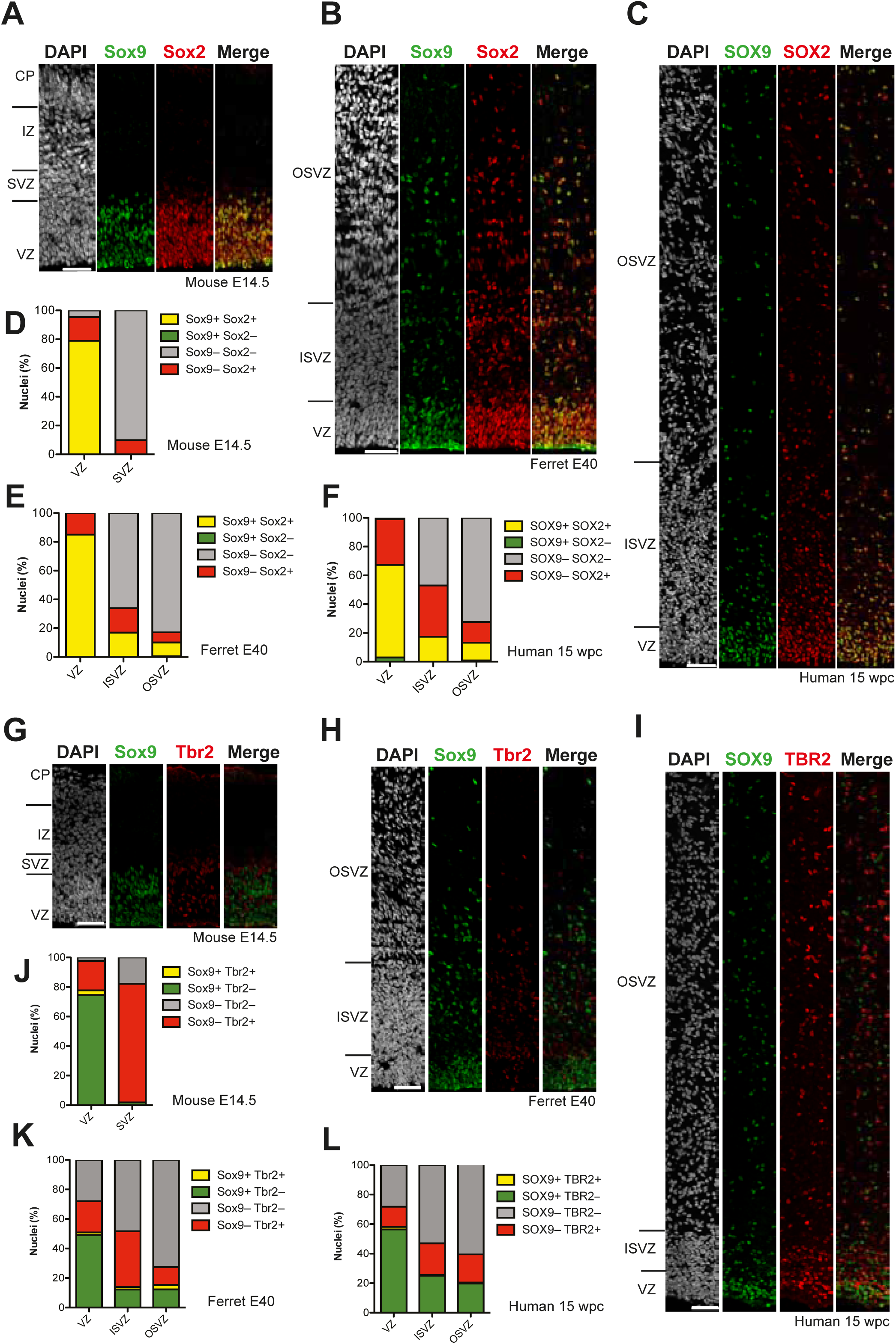
Sox9-expressing BPs occur in the SVZ of embryonic ferret and fetal human but not embryonic mouse neocortex. (**A-C**) Double immunofluorescence for Sox9 (green) and Sox2 (red), combined with DAPI staining (white), of mouse E14.5 (**A**), ferret E40 (**B**) and human 15 wpc (**C**) neocortex. (**D-F**) Quantification of the percentage of nuclei (identified by DAPI staining) that are Sox9 plus Sox2 double-positive (yellow), Sox9-positive only (green), Sox2-positive only (red), and Sox9 plus Sox2 double-negative (grey), in mouse E14.5 (**D**), ferret E40 (**E**) and human 15 wpc (**F**) neocortex. (**G-I**) Double immunofluorescence for Sox9 (green) and Tbr2 (red), combined with DAPI staining (white), of mouse E14.5 (**G**), ferret E40 (**H**) and human 15 wpc (**I**) neocortex. (**J-L**) Quantification of the percentage of nuclei (identified by DAPI staining) that are Sox9 plus Tbr2 double-positive (yellow), Sox9-positive only (green), Tbr2-positive only (red), and Sox9 plus Tbr2 double-negative (grey), in mouse E14.5 (**J**), ferret E40 (**K**) and human 15 wpc (**L**) neocortex. (**A-C, G-I**) Ventricular surface is down. Upper margins of images in (**A, B, G, H**) correspond to the pial surface (**A, G**) and the basal boundary of the OSVZ (**B, H**); in (**C, I**) most but not all of the OSVZ is shown due to space constraints. Scale bars, 50 µm.

In light of these mRNA data, we analyzed the expression of the Sox9/SOX9 protein in developing mouse and human neocortex by immunofluorescence. For these analyses, we included a third species, the ferret, which like human but in contrast to mouse exhibits an expanded SVZ containing BPs with proliferative capacity (Borrell and Reillo, 2012; Gertz et al., 2014; Kawasaki, 2018; LaMonica et al., 2013; Reillo et al., 2011; Turrero Garcia et al., 2015). Consistent with the *Sox9/SOX9* mRNA data (see Supp Fig. 1), the Sox9/SOX9 protein was restricted to the VZ in E14.5 mouse neocortex (Fig. 1A, G), but was abundantly expressed in the VZ, ISVZ and OSVZ of E40 ferret and 15 wpc human neocortex (Fig. 1B, C, H, I). At the subcellular level, the Sox9/SOX9 protein showed a nuclear localization. These data suggested that the Sox9/SOX9 protein is expressed in BPs of ferret and human, but not mouse, neocortex.

In order to further explore the identity of the Sox9-expressing cells in the SVZ of developing ferret and human neocortex, we investigated a potential co-expression of Sox9 with Sox2, the expression of which among BPs is characteristic of proliferative BPs, notably bRG (Graham et al., 2003; Hansen et al., 2010; Reillo et al., 2011; Wang et al., 2011). Double immunofluorescence showed that virtually all Sox9-expressing cells in the ISVZ and OSVZ of E40 ferret and 15 wpc human neocortex co-expressed Sox2, implying that these cells were BPs, and that these cells comprised about half of the Sox2-positive BPs (Fig. 1B, C, E, F). Of note, regarding the VZ of developing mouse, ferret and human neocortex, while nearly all cells were Sox2-positive, 70-80% of these also expressed Sox9 (Fig. 1A-F), again in line with these cells being cNPCs.

Analysis of expression of the transcription factor Tbr2 (encoded by *Eomes*) allows one to identify newborn bIPs in the VZ, and can aid in distinguishing between bIPs and bRG in the SVZ (Englund et al., 2005; Kowalczyk et al., 2009; Pontious et al., 2008; Sessa et al., 2008). Double immunofluorescence of the E14.5 mouse, E40 ferret and 15 wpc human neocortex for Sox9 and Tbr2 revealed that in all germinal zones, virtually none of the Sox9-positive cells expressed Tbr2, indicative of a mutually exclusive expression pattern (Fig. 1G-L). Regarding the VZ, these data imply that the Sox9 and Sox2 double-positive cells observed in the three species (Fig. 1D-F) are aRG and the Sox9-negative but Sox2-positive cells (Fig. 1D-F) are newborn BPs, notably newborn bIPs. Regarding the SVZ, these data suggest that the Sox9 and Sox2 double-positive cells observed in ferret and human are bRG rather than neurogenic bIPs. The latter conclusion in turn implies that Sox9 expression is down-regulated upon BPs becoming committed to neurogenesis.

### Sox9-expressing BPs in ferret and human can re-enter the cell cycle and include bRG

The data described so far prompted us to further study the Sox9-expressing cNPCs, notably the BPs, and to analyze their proliferative capacity. For this purpose, we used postnatal ferret kits, which allow application of a broad spectrum of experimental approaches (Fietz et al., 2010; Gertz et al., 2014; Kawasaki, 2018; Reillo and Borrell, 2012; Turrero Garcia et al., 2015). We first performed immunofluorescence for the cell cycle marker PCNA on postnatal day (P) 2 (Fig. 2A) and P3 ferret neocortex and quantified the percentage of Sox9-positive cells that were cycling. This was found to be the case for 91% and 81% of them in the VZ, for 67% and 75% in the ISVZ, and for 87% and 94% in the OSVZ at P2 and P3, respectively (Fig. 2B, C).

**Fig. 2.**
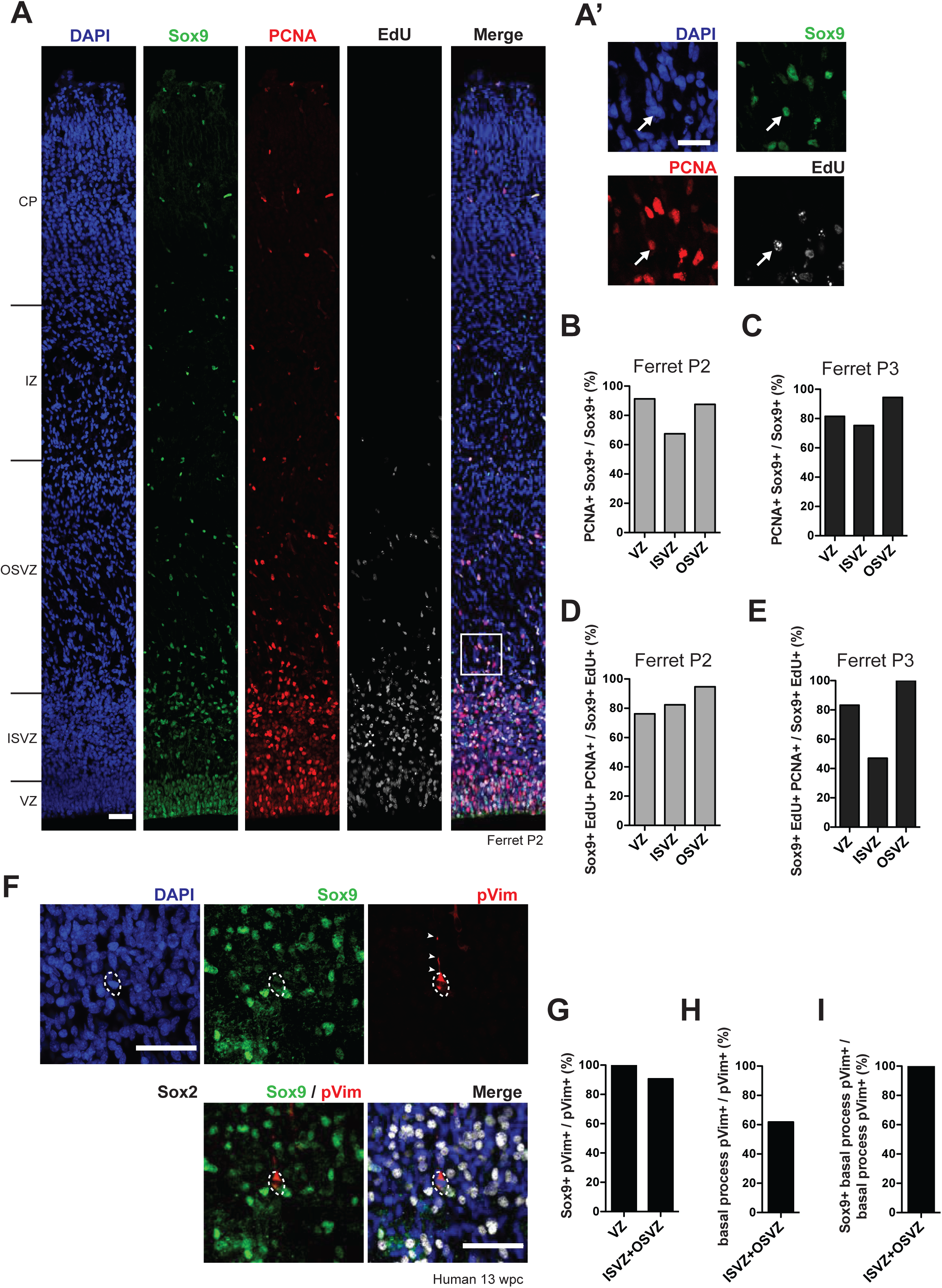
Sox9-expressing BPs in the SVZ of developing ferret and human neocortex are capable of cell cycle re-entry and include bRG. **(A)** Triple (immuno)fluorescence for Sox9 (green), PCNA (red) and EdU (white), combined with DAPI staining (blue), of P2 ferret neocortex. EdU was administrated at P0. Boxed area in (**A**) is shown in higher magnification in **(A’)**. White arrows indicate a nucleus that is triple-positive for Sox9, PCNA and EdU, i.e. a Sox9-positive cycling BP in the OSVZ. **(B, C)** Quantification of the percentage of Sox9-positive nuclei that are PCNA-positive, i.e. the percentage of Sox9+ cells that are cycling, in the indicated germinal zones of P2 (**B**) and P3 (**C**) ferret neocortex. **(D, E)** Quantification of the percentage of Sox9 and EdU double-positive nuclei that are PCNA-positive, i.e. the percentage of Sox9+ cNPCs that have re-entered the cell-cycle, in the indicated germinal zones of P2 (**D**) and P3 (**E**) ferret neocortex. EdU was administrated at P0. **(F)** Triple immunofluorescence for Sox9 (green), phospho-vimentin (pVim, red) and Sox2 (white), combined with DAPI staining (blue), in the OSVZ of 13 wpc human neocortex. Arrowheads indicate the basal process, and dashed lines delineate the mitotic cell body, of a bRG. **(G)** Quantification of the percentage of pVim-positive cells, i.e. of mitotic cNPCs, that are Sox9-positive, in the indicated germinal zones of 13 wpc human neocortex. **(H)** Quantification of the percentage of pVim-positive cells that bear a basal process, i.e. the percentage of mitotic BPs that are bRG, in the ISVZ plus OSVZ of 13 wpc human neocortex. **(I)** Quantification of the percentage of basal process-bearing pVim-positive cells, i.e. of mitotic bRG, that are Sox9-positive, in the ISVZ plus OSVZ of 13 wpc human neocortex. (**A, A’, F**) Scale bars, 50 µm (**A, F**) and 25 µm (**A’**). In (**A**), ventricular surface is down.

We next determined if the Sox9-positive cNPCs were capable of cell cycle re-entry. To this end, we administered the thymidine analog EdU to P0 ferret kits in order to label cells in S-phase, and sacrificed the kits at P2 and P3, that is, after a time interval that – in light of the known cell cycle length of ferret postnatal cNPCs (Turrero Garcia et al., 2015) – should allow the labeled cells to go through mitosis and the resulting daughter cells to become either post-mitotic or to re-enter the cell cycle. Cell cycle re-entry was assessed by immunofluorescence for PCNA of Sox9 and EdU double-positive cells (Fig. 2A and A’). At P2, 76% of the Sox9-positive progeny in the VZ, 82% in the ISVZ and 94% in the OSVZ had re-entered the cell cycle (Fig. 2D). Similar values were observed for the VZ and OSVZ at P3 (Fig. 2E), suggesting that even the granddaughter cells of the EdU-labeled cNPCs maintained proliferative capacity. In contrast, for the ISVZ, the value decreased to 47%, in line with the progression of neural development, specifically the reduction in proliferative capacity and the increased generation of neurons in this germinal zone (Reillo et al., 2011; Turrero Garcia et al., 2015) (Fig. 2E). We conclude that in the developing ferret neocortex, the Sox9-expressing BPs in the OSVZ, like the Sox9-expressing aRG in the VZ, exhibit a high proliferative capacity.

We sought to extend this investigation to fetal human neocortex and to focus specifically on bRG, which constitute a major BP subpopulation in the human SVZ (Fietz et al., 2010; Hansen et al., 2010). To identify Sox9-expressing bRG, we combined immunofluorescence for Sox9 with that for phospho-vimentin, which labels cNPCs in mitosis and – by staining the basal process – marks mitotic bRG (Fietz et al., 2010; Florio et al., 2015; Hansen et al., 2010) (Fig. 2F). In 13 wpc human neocortex, essentially all mitotic aRG in the VZ and the overwhelming majority (91%) of the mitotic BPs in the ISVZ plus OSVZ were Sox9-positive (Fig. 2G). Moreover, in line with previous observations (Fietz et al., 2010), 61% of the mitotic BPs exhibited a basal process and thus were bRG (Fig. 2H). Importantly, all of the latter expressed Sox9, indicating that bRG in the fetal human neocortex are Sox9-positive (Fig. 2I).

### Conditional Sox9 expression in mouse BPs increases their proliferation and cell cycle re-entry

The absence of Sox9 expression in mouse BPs, which are known to lack proliferative capacity, vs. the presence of Sox9 expression in ferret/human BPs, which exhibit proliferative capacity (Fietz and Huttner, 2011; Florio and Huttner, 2014; Lui et al., 2011; Reillo and Borrell, 2012), led us to hypothesize that Sox9 could have a role in promoting a proliferative state of BPs. We therefore sought to conditionally express Sox9 in mouse BPs (in addition to its physiological expression in aRG), to see if this would promote mouse BP proliferation. To this end, we generated a conditional Sox9 expression construct (Fig. 3A) and introduced it by *in utero* electroporation into aRG of tamoxifen-treated E13.5 embryos of the *Tis21*-CreER^T2^ mouse line, which allows expression of floxed constructs specifically in BP-genic aRG and the BP progeny derived therefrom (Wong et al., 2015) (Fig. 3B). When using the conditional Sox9 expression construct in conjunction with tamoxifen-induced Cre-mediated recombination, expression of nuclear RFP is indicative of Sox9 expression (Fig. 3A).

**Fig. 3.**
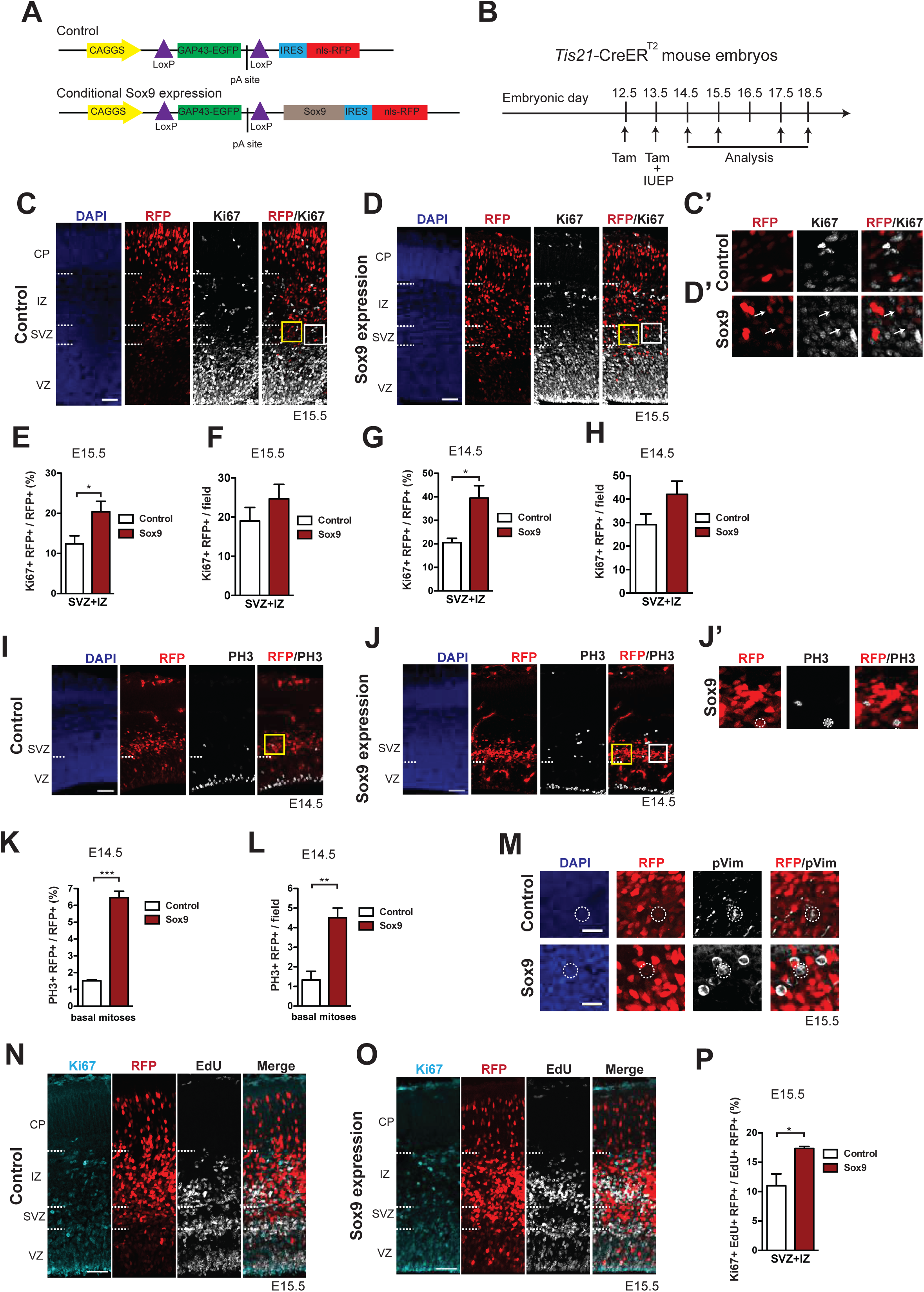
Conditional Sox9 expression in BPs of embryonic mouse neocortex increases their proliferation and cell cycle re-entry. **(A)** Constructs used to conditionally express nuclear RFP without (top, control construct) and with (bottom, conditional Sox9 expression construct) Sox9 in mouse BPs and their progeny using the *Tis21*-CreER^T2^ line (see **B**). **(B)** Workflow of tamoxifen administration (Tam) at E12.5 and E13.5, *in utero* electroporation (IUEP) at E13.5, and immunostaining analyses of the neocortex at the indicated time points (arrows) yielding the data shown in this figure and subsequent figures, using heterozygous *Tis21*-CreER^T2^ mouse embryos. **(C, D)** Double immunofluorescence of neocortex for RFP (red) and Ki67 (white), combined with DAPI-staining (blue), 48 hours after electroporation of control construct (**C**) or conditional Sox9 expression construct (**D**). Dashed lines indicate the borders between VZ, SVZ, IP and CP. White boxed areas of the SVZ in (**C**) and (**D**) are shown at higher magnification in (**C’**) and (**D’**), respectively; arrows indicate RFP-positive nuclei that are Ki67-positive. Yellow boxed areas of the SVZ in (**C**) and (**D**) are shown at higher magnification in Fig. 6A, top row and bottom row, respectively. **(E-H)** Quantifications in the neocortical SVZ plus IZ, upon electroporation of control construct (white columns) or conditional Sox9 expression construct (red columns). **(E)** Quantification of the percentage of RFP-positive nuclei that are Ki67-positive, 48 hours after electroporation. **(F)** Quantification of the number of Ki67 and RFP double-positive nuclei per microscopic field of 200-µm apical width, 48 hours after electroporation. **(G)** Quantification of the percentage of RFP-positive nuclei that are Ki67-positive, 24 hours after electroporation. Related representative images are not shown. **(H)** Quantification of the number of Ki67 and RFP double-positive nuclei per microscopic field of 200-µm apical width, 24 hours after electroporation. Related representative images are not shown. **(I, J)** Double immunofluorescence of neocortex for RFP (red) and phosphohistone H3 (PH3, white), combined with DAPI staining (blue), 24 hours after electroporation of control construct **(I)** or conditional Sox9 expression construct **(J)**. Dashed lines indicate the border between VZ and SVZ. White boxed area in (**J**) is shown at higher magnification in (**J’**); dashed circles delineate a basal, PH3 and RFP double-positive mitosis. Yellow boxed area of the SVZ in (**J**) is shown at higher magnification in Fig. 6B. **(K)** Quantification of the percentage of abventricular RFP-positive cells in neocortex that undergo basal mitosis as revealed by PH3 immunofluorescence, 24 hours after electroporation of control construct (white column) or conditional Sox9 expression construct (red column). **(L)** Quantification of the number of basal, PH3 and RFP double-positive mitoses in neocortex per microscopic field of 200-µm apical width, 24 hours after electroporation of control construct (white column) or conditional Sox9 expression construct (red column). **(M)** Double immunofluorescence of neocortex for RFP (red) and phospho-vimentin (pVim, white), combined with DAPI staining (blue), of the SVZ, 48 hours after electroporation of control construct (white column) or conditional Sox9 expression construct (red column). Dashed circles delineate the cell body of RFP and pVim double-positive cells. **(N, O)** Triple (immuno)fluorescence of neocortex for Ki67 (cyan), RFP (red) and EdU (white) 48 hours after electroporation of control construct **(N)** or conditional Sox9 expression construct **(O)**. A single pulse of EdU was administered at E14.5, i.e. 24 hours after electroporation and 24 hours prior to analysis. Dashed lines indicate the borders between VZ, SVZ, IZ and CP. **(P)** Quantification of the percentage of RFP and EdU double-positive nuclei in the neocortical SVZ plus IZ that are Ki67-positive, i.e. the percentage of RFP+ BPs that have re-entered the cell-cycle, 48 hours after electroporation of control construct (white column) or conditional Sox9 expression construct (red column) and 24 hours after EdU administration at E14.5. **(C, D, I, J, M, N, O)** Scale bars, 50 *μ*m **(C, D, I, J, N, O)**, 20 *μ*m **(M)**. **(E, F, G, H, K, L, P)** Two-tailed, unpaired *t*-test: * p<0.05, ** p<0.01, *** p<0.001. Data are the mean of 6 **(E, F)**, 4 **(G, H)** and 3 **(K, L, P)** embryos, each from a different litter; for each embryo, two microscopic fields, each of 200-µm apical width, were counted, and the values obtained were averaged. Error bars represent SEM.

We first validated the conditional Sox9 expression construct by transfecting it into HEK293T cells, which do not express endogenous Sox9 (Supp Fig. 2A, B), with or without co-transfection of a construct driving constitutive expression of nuclear Cre. Transfection of the conditional Sox9 expression construct alone led to the expression of GFP from a floxed cassette, but not of RFP, the expression of which depends on the removal of the floxed GFP cassette (Supp Fig. 2C, see also Fig. 3A). In contrast, transfection of the conditional Sox9 expression construct in combination with the Cre-expressing construct resulted in the expression of RFP but not GFP, and all RFP-expressing cells were positive for Sox9 (Supp Fig. 2D). These observations provided a validation of the conditional Sox9 expression construct. It is worth noting that whereas in these transfected HEK293T cells the RFP was diffusely distributed in the nucleoplasm, Sox9 immunoreactivity was concentrated at the inner surface of the nuclear envelope (Supp Fig. 2D).

Next, we validated the conditional expression of Sox9 in mouse BP-genic aRG and their BP progeny using the *Tis21*-CreER^T2^ line. We induced translocation of Cre to the nucleus by tamoxifen administration at E12.5 and E13.5, introduced either the control RFP-expressing construct (see Fig. 3A) or the conditional Sox9 expression construct into neocortical aRG by *in utero* electroporation at E13.5, and performed immunofluorescence analyses at E14.5 (see Fig. 3B). Upon electroporation of the control construct, we observed expression of RFP predominantly in the SVZ, whereas the expression of the endogenous Sox9 was confined to the VZ (Supp Fig. 3A), consistent with the data described above (see Fig. 1A). In contrast, upon electroporation of the conditional Sox9 expression construct, strong Sox9 immunoreactivity was observed not only in the VZ, but also in the SVZ (Supp Fig. 3B). Importantly, all of the strongly Sox9-positive, i.e. exogenous Sox9-expressing, cells co-expressed nuclear RFP (Supp Fig. 3B), demonstrating that RFP expression from the conditional Sox9 expression construct can be taken as an indicator of Sox9 expression. Furthermore, these data show that the use of the conditional Sox9 expression construct in tamoxifen-treated *Tis21*-CreER^T2^ embryos is a means of eliciting Sox9 expression in mouse BPs.

These findings provided a basis to investigate the effects of conditional Sox9 expression in mouse BPs on their proliferative capacity. We first examined the expression of Ki67, a marker of cycling cells, upon electroporation of the mouse neocortex at E13.5 (Fig. 3B). Indeed, conditional Sox9 expression doubled the proportion of cells in the SVZ and intermediate zone (IZ) derived from the targeted cells (as revealed by RFP expression) that were cycling and hence were BPs, both upon analysis at E15.5 (Fig. 3C-E) and at E14.5 (Fig. 3G). Conditional Sox9 expression also slightly increased the absolute number of cycling, targeted cell-derived (RFP+) BPs per unit area, both at 24 hours and 48 hours after electroporation, although this effect was not statistically significant (Fig. 3F and H).

To corroborate the effect of conditional Sox9 expression on mouse BP proliferation, we analyzed the abundance of basal mitoses by performing immunofluorescence for phosphohistone H3 one day after electroporation at E13.5 (Fig. 3I, J, J’). Conditional Sox9 expression markedly increased the proportion of the targeted cell-derived (RFP+) BPs that underwent mitosis (Fig. 3K) and the absolute number of targeted cell-derived (RFP+) mitotic BPs per unit area (Fig. 3L).

In mouse, the overwhelming majority of BPs in the embryonic lateral neocortex at mid-neurogenesis are bIPs, with bRG constituting only a minor fraction (Arai et al., 2011; Florio and Huttner, 2014; Shitamukai et al., 2011; Vaid et al., 2018; Wang et al., 2011). Immunofluorescence for phospho-vimentin at E15.5 did not provide evidence that a noteworthy fraction of the targeted cell-derived (RFP+) mitotic BPs observed upon conditional Sox9 expression exhibited a basal process (Fig. 3M). This suggested that the Sox9-induced increase in the proliferation of mouse BPs pertained primarily to bIPs.

We next investigated whether the increase in mouse BPs upon conditional Sox9 expression was accompanied by an increase in cell cycle re-entry. To this end, we analyzed Ki67 immunofluorescence at E15.5 of mouse neocortex electroporated at E13.5 and subjected to EdU pulse-labeling at E14.5 (Fig. 3N, O). This revealed that conditional Sox9 expression significantly increased the proportion of targeted cell-derived (RFP+), EdU-containing cells in the SVZ plus IZ that were Ki67+ and hence were cycling BPs (Fig. 3P). These data are consistent with the progeny of BPs exhibiting an increased ability to re-enter the cell cycle, i.e. to remain being BPs (as opposed to becoming postmitotic neurons).

### Conditional Sox9 expression in mouse BPs reduces Tbr2 expression and induces premature gliogenesis

BPs in the E13.5-15.5 mouse neocortex, i.e. bIPs, are typically neurogenic, undergoing symmetric consumptive division that generates two neurons (Haubensak et al., 2004; Miyata et al., 2004; Noctor et al., 2004), and express the transcription factor marker Tbr2 (Fig. 1G, J) (Englund et al., 2005; Kowalczyk et al., 2009; Pontious et al., 2008; Sessa et al., 2008). We asked whether conditional Sox9 expression, in addition to increasing the proliferative capacity of mouse BPs, would alter their identity and fate. Tbr2 immunofluorescence of mouse neocortex electroporated at E13.5 revealed that conditional Sox9 expression decreased the abundance of Tbr2-positive cells at E15.5 (Fig. 4A, B), especially in the SVZ (Fig. 4A’, B’). Specifically, conditional Sox9 expression reduced the proportion of targeted cell-derived (RFP+) BPs in the SVZ that expressed Tbr2 to half (Fig. 4C), and markedly decreased their abundance per unit area (Fig. 4D). These findings provided a first indication that conditional Sox9 expression in mouse BPs may alter their identity and fate, possibly reducing their commitment to neurogenesis.

**Fig. 4.**
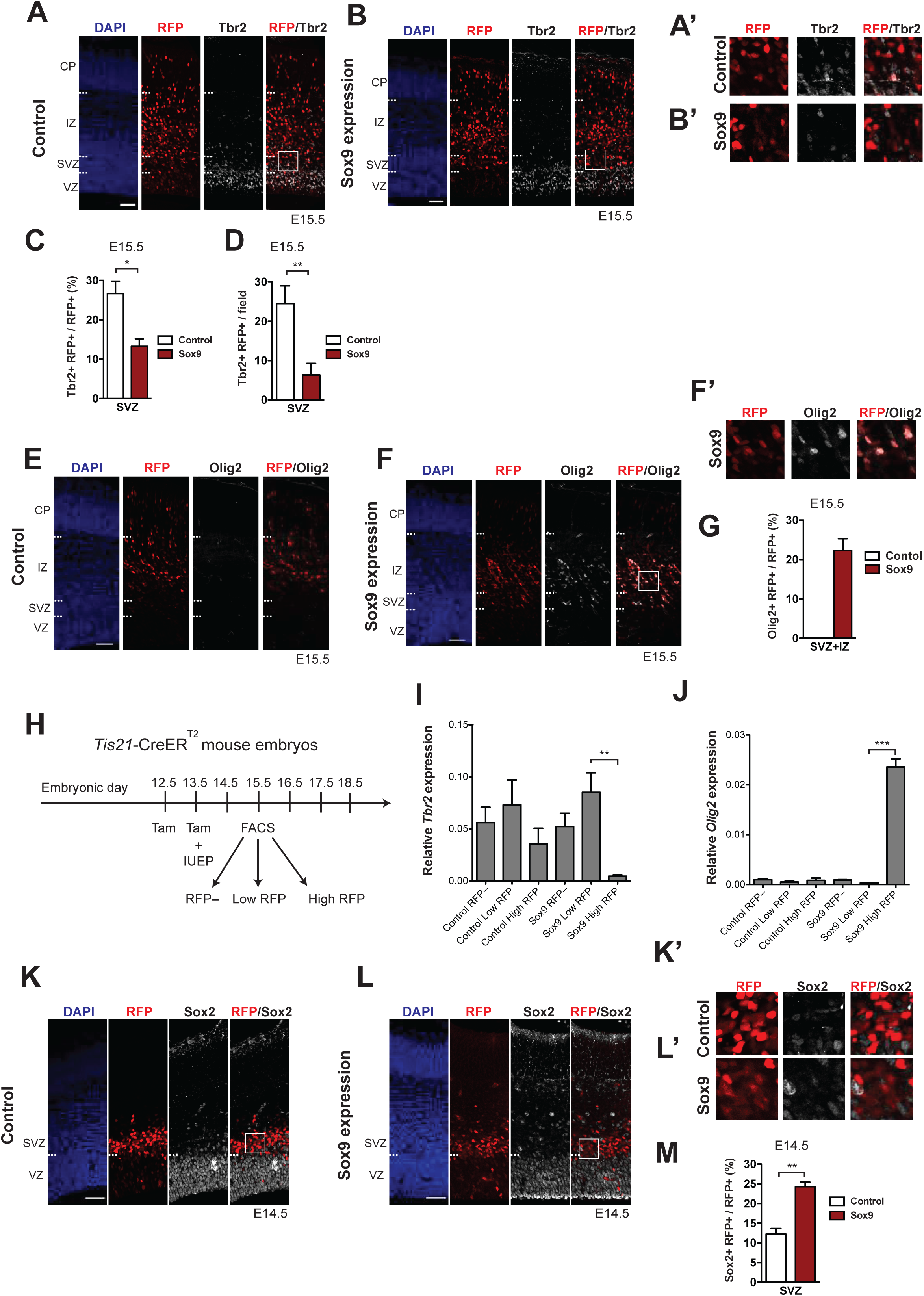
Conditional Sox9 expression in mouse BPs represses Tbr2 expression and induces premature gliogenesis in a dose-dependent manner. Heterozygous *Tis21*-CreER^T2^ mouse embryos received tamoxifen administration at E12.5 and E13.5 and were subjected to *in utero* electroporation of the neocortex at E13.5 with either control construct or conditional Sox9 expression construct, followed by immunostaining analyses either 24 hours (**K-M**) or 48 hours (**A-G**) later. **(A, B)** Double immunofluorescence for RFP (red) and Tbr2 (white), combined with DAPI staining (blue), 48 hours after electroporation of control construct (**A**) or conditional Sox9 expression construct (**B**) (see Fig. 3A). Dashed lines indicate the borders between VZ, SVZ, IP and CP. Boxed areas of the SVZ in (**A**) and (**B**) are shown at higher magnification in (**A’**) and (**B’**) respectively. **(C)** Quantification of the percentage of RFP-positive nuclei in the SVZ that are Tbr2-positive, 48 hours after electroporation of control construct (white column) or conditional Sox9 expression construct (red column). **(D)** Quantification of the number of Tbr2 and RFP double-positive nuclei in the SVZ per microscopic field of 200-µm apical width, 48 hours after electroporation of control construct (white column) or conditional Sox9 expression construct (red column). **(E, F)** Double immunofluorescence for RFP (red) and Olig2 (white), combined with DAPI staining (blue), 48 hours after electroporation of control construct (**E**) or conditional Sox9 expression construct (**F**). Dashed lines indicate the borders between VZ, SVZ, IP and CP. Boxed area of IZ in **(F)** is shown in higher magnification in **(F’)**. **(G)** Quantification of the percentage of RFP-positive nuclei in the SVZ plus IZ that are Olig2-positive, 48 hours after electroporation of control construct (white column) or conditional Sox9 expression construct (red column). **(H)** Workflow of tamoxifen administration (Tam) at E12.5 and E13.5, *in utero* electroporation (IUEP) of the neocortex at E13.5, and FACS at E15.5 followed by qPCR analyses of RFP-positive cells yielding the data shown in **(I)** and **(J)**, using heterozygous *Tis21*-CreER^T2^ mouse embryos. **(I, J)** Quantification of *Tbr2* **(I)** and *Olig2* **(J)** mRNA levels relative to the *Gapdh* mRNA level by qPCR analysis, in RFP-negative, low-level RFP-expressing and high-level RFP-expressing cell populations isolated by FACS (see **H**) 48 hours after electroporation of control construct (left three columns) or conditional Sox9 expression construct (right three columns). **(K, L)** Double immunofluorescence for RFP (red) and Sox2 (white), combined with DAPI staining (blue), 24 hours after electroporation of control construct (**K**) or conditional Sox9 expression construct (**L**). Dashed lines indicate border between VZ and SVZ. Boxed areas in SVZ in (**K**) and (**L**) are shown at higher magnification in **(K’)** and **(L’)**, respectively. **(M)** Quantification of the percentage of RFP-positive nuclei in the SVZ that are Sox2-positive, 24 hours after electroporation of control construct (white column) or conditional Sox9 expression construct (red column). **(A, B, E, F, K, L)** Scale bars, 50 µm. **(C, D, I, J, M)** Two-tailed, unpaired *t*-test: * p<0.05, ** p<0.01, *** p<0.001. Data are the mean of 3 **(C, D, G, M)**, 5 **(I)** and 4 **(J)** embryos, each from a different litter; for each embryo in **(C, D, G, M)**, two microscopic fields, each of 200-µm apical width, were counted, and the values obtained were averaged. Error bars represent SEM.

In light of the involvement of Sox9 in the switch of NPCs from neurogenesis to gliogenesis reported for the developing mouse cerebellum and spinal cord (Finzsch et al., 2008; Kang et al., 2012; Molofsky et al., 2013; Stolt et al., 2003; Stolt and Wegner, 2010; Vong et al., 2015; Wegner and Stolt, 2005), we explored a potential role of Sox9 in gliogenesis in the developing neocortex. We first examined the Sox9-positive BPs in the ISVZ and OSVZ of the developing ferret neocortex (see Fig. 1B, E, H, K) for the expression of Olig2, a transcription factor implicated in gliogenesis (Marshall et al., 2005; Rowitch and Kriegstein, 2010; Zhou et al., 2001). Double immunofluorescence for Sox9 and Olig2 of E40 and P1 ferret neocortex revealed scattered double-positive cells in the ferret ISVZ and OSVZ (see Supp Fig. 4A for E40), which upon quantification amounted to 14% and 22% of the Sox9-positive cells in these germinal zones at E40 and P1, respectively (Supp Fig. 4B). Given that essentially all Sox9-positive cells in the E40 (Fig. 1E, Supp Fig. 4C) and P1 (Supp Fig. 4C) ISVZ and OSVZ of ferret neocortex are Sox2-positive and hence BPs, we conclude that up to a quarter of these progenitors are committed to gliogenesis at the developmental stages studied.

We next examined whether mouse BPs conditionally expressing Sox9 include gliogenic precursor cells. No Olig2 immunoreactivity was detected in the E15.5 neocortical wall by immunofluorescence upon control electroporation at E13.5 (Fig. 4E), in line with the onset of neocortical gliogenesis in mouse occurring later, i.e. around E18.5 (Miller and Gauthier, 2007). Remarkably, however, upon conditional Sox9 expression for two days, many Olig2-positive cells were observed in the SVZ and IZ (Fig. 4F, F’). Quantification of Olig2 and RFP double-positive cells revealed that 22% of the targeted cell-derived (RFP+) BPs in the mouse E15.5 SVZ and IZ had adopted a gliogenic identity (Fig. 4G).

We examined whether Olig2 expression was related to the level of Sox9 expression. To this end, we performed *in utero* electroporation of the neocortex of tamoxifen-treated E13.5 *Tis21*-CreER^T2^ mouse embryos with either the control RFP-expressing construct or the conditional Sox9 expression construct, dissociated the neocortical cells at E15.5, and isolated RFP+ cells by FACS (Fig. 4H). With this approach, we obtained three cell populations each from control- and Sox9-electroporated neocortex, referred to as RFP–, Low RFP and High RFP (Supp Fig. 5A, B). We first examined *Sox9* mRNA levels by quantitative PCR on these sorted cell populations and observed that relative *Sox9* expression was highest in the High RFP population from Sox9-electroporated neocortex and was much lower in the Low RFP population from the same sample (Supp Fig. 5C). The other sorted cell populations (RFP–, Low RFP and High RFP of control, RFP– from Sox9-electroporated neocortex) showed almost no *Sox9* expression (Supp Fig. 5C). We next analyzed mRNA levels of *Tbr2* and *Olig2* and found that relative expression of *Tbr2* was almost completely repressed (Fig. 4I) and relative expression of *Olig2* was specifically induced (Fig. 4J) in the High RFP population of Sox9-electroporated neocortex. In contrast, the other sorted cell populations showed *Tbr2* expression but no detectable *Olig2* expression (Fig. 4I and J). We conclude that high Sox9 levels in mouse BPs induce a switch in their cell fate to gliogenesis by inducing *Olig2* and repressing *Tbr2* expression.

**Fig. 5.**
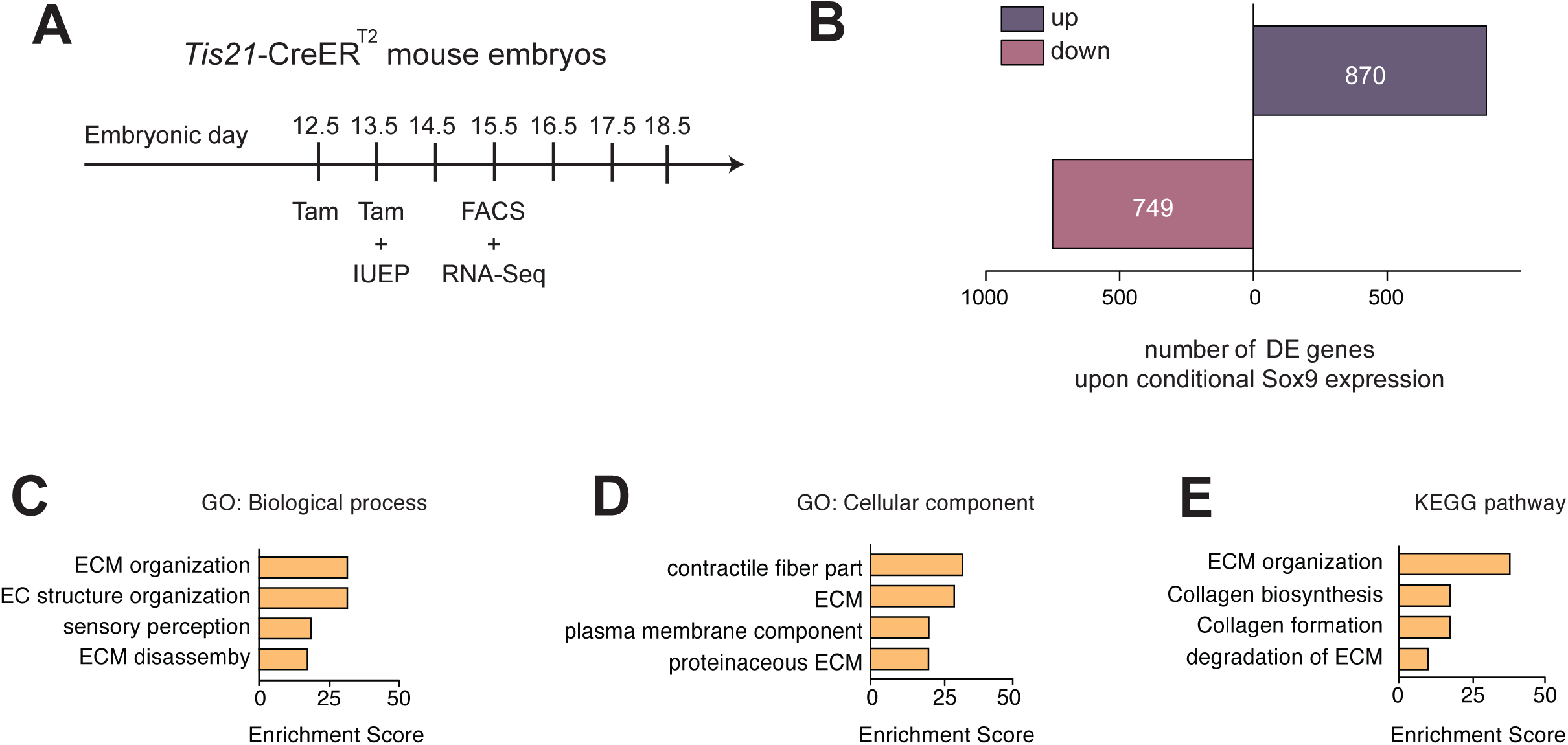
mRNAs upregulated upon conditional Sox9 expression in mouse BPs are mainly related to ECM. **(A)** Workflow of tamoxifen administration (Tam) at E12.5 and E13.5, *in utero* electroporation (IUEP) of the neocortex at E13.5 with either control construct or conditional Sox9 expression construct, and FACS at E15.5 followed by RNA-Seq analysis of high-level RFP-expressing cells yielding the data shown in (**B**-**E**), using heterozygous *Tis21*-CreER^T2^ mouse embryos. **(B)** Number of mRNAs of protein-encoding genes differentially expressed (DE) 48 hours after electroporation of conditional Sox9 expression construct as compared to control construct (see Fig. 3A); up, upregulated; down, downregulated. **(C-E)** Gene ontology (GO) term enrichment analyses for biological process **(C)** and cellular component **(D)**, and KEGG pathway analysis **(E)**, using as input the set of 870 genes upregulated upon conditional Sox9 expression. The top 4 enriched terms/pathways (p<0.01) are shown.

The Sox9-induced, dose-dependent switch of BPs to a gliogenic fate raised the question whether all BPs had lost their neuronal progenitor identity. To answer this question, we immunostained control- and Sox9-electroporated neocortex for Sox2, at E14.5 (Fig. 4K and L). Specifically, we examined the expression of Sox2 in the SVZ (Fig. 4K’ and L’). Quantification of the percentage of the RFP+ cells in the SVZ that were Sox2-positive revealed an increase upon Sox9 expression (Fig. 4M). Given that upon conditional Sox9 expression at E13.5, 10-15% of the RFP+ cells in the SVZ are still positive for Tbr2 at E15.5 (Fig. 4C), this observation suggests that the increased number of BPs consist of a mix of neurogenic and gliogenic progenitors.

### Conditional Sox9 expression in mouse BPs mainly upregulates transcription of ECM components

In light of the findings described so far, it was important to obtain a global view of the transcriptional effects of conditional Sox9 expression in mouse BPs. To this end, we compared their transcriptomes upon electroporation of tamoxifen-treated E13.5 *Tis21*-CreER^T2^ mouse neocortex with either the control or the conditional Sox9 expression construct, followed by dissociation of the electroporated tissue at E15.5, isolation of the High RFP cells by FACS (Supp Fig. 5), and RNA sequencing (Fig. 5A). We first assessed the difference between control- and Sox9-electroporated samples by principal component analysis (PCA), which revealed the two groups of samples to be distinct (Supp Fig. 6A). We then assembled a sample distance matrix of all expressed genes and observed that the conditionally Sox9 expressing samples clustered separately from the control samples (Supp Fig. 6B).

**Fig. 6.**
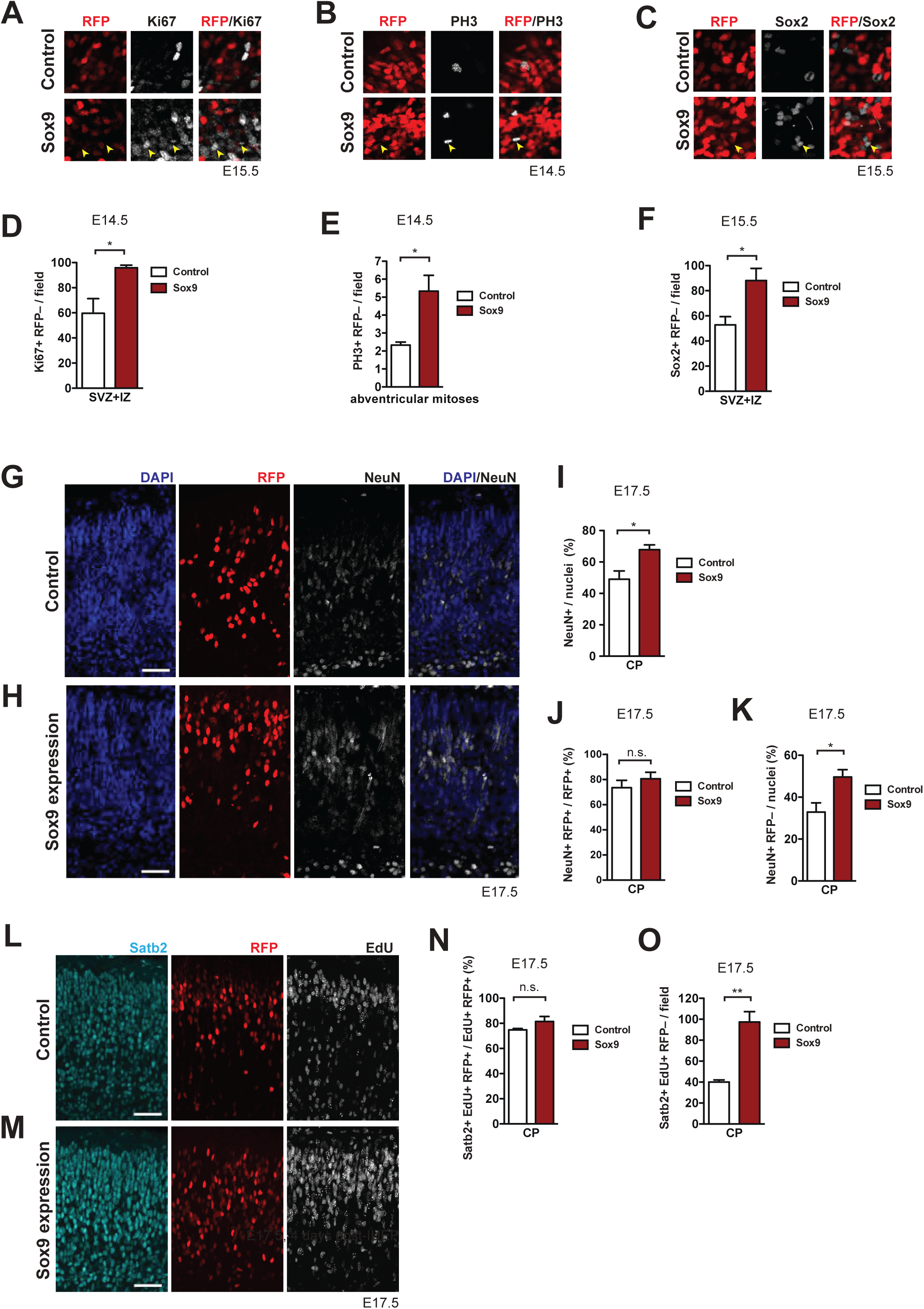
Conditional Sox9 expression in mouse BPs induces cell non-autonomous proliferation and neuron production. Heterozygous *Tis21*-CreER^T2^ mouse embryos received tamoxifen at E12.5 and E13.5, were subjected to *in utero* electroporation of the neocortex with either control construct or conditional Sox9 expression construct (see Fig. 3A) at E13.5, and subjected to immunostaining analyses of the neocortex at the time points indicated in (**A**-**O**). **(A-C)** High-magnification images of the SVZ showing double immunofluorescence for Ki67 (white) and RFP (red) (see yellow boxed areas in Fig. 3C, **D**) **(A)**, phosphohistone H3 (PH3, white) and RFP (red) (see yellow boxed area in Fig. 3I, J) **(B)**, and Sox2 (white) and RFP (red), 48 hours after electroporation of control or conditional Sox9 expression construct **(C)**. Yellow arrowheads indicate BPs that are negative for RFP, but positive for Ki67 **(A)**, PH3 **(B)** or Sox2 **(C)**. **(D)** Quantification of the number of Ki67-positive, RFP-negative nuclei in the SVZ plus IZ per microscopic field of 200-µm apical width, 24 hours after electroporation of control construct (white column) or conditional Sox9 expression construct (red column). **(E)** Quantification of the number of abventricular, PH3-positive, RFP-negative mitoses per microscopic field of 200-µm apical width, 24 hours after electroporation of control construct (white column) or conditional Sox9 expression construct (red column). **(F)** Quantification of the number of Sox2-positive, RFP-negative nuclei in the SVZ plus IZ per microscopic field of 200-µm apical width, 48 hours after electroporation of control construct (white column) or conditional Sox9 expression construct (red column). **(G, H)** Double immunofluorescence for RFP (red) and NeuN (white), combined with DAPI staining (blue), in the CP 4 days after electroporation of control construct (**G**) or conditional Sox9 expression construct (**H**). **(I)** Quantification of the percentage of nuclei (identified by DAPI staining) in the CP that are NeuN-positive, 4 days after electroporation of control construct (white column) or conditional Sox9 expression construct (red column). **(J)** Quantification of the percentage of RFP-positive nuclei in the CP that are NeuN-positive, 4 days after electroporation of control construct (white column) or conditional Sox9 expression construct (red column). **(K)** Quantification of the percentage of nuclei (identified by DAPI staining) in the CP that are NeuN-positive and RFP-negative, 4 days after electroporation of control construct (white column) or conditional Sox9 expression construct (red column). **(L, M)** Triple (immuno)staining for Satb2 (cyan), RFP (red) and EdU (white) in the CP, 4 days after electroporation of control construct (**L**) or conditional Sox9 expression construct (**M**). A single pulse of EdU was administered at E14.5, i.e. 24 hours after electroporation and 3 days prior to analysis. **(N)** Quantification of the percentage of EdU and RFP double-positive nuclei in the CP that are Satb2-positive, i.e. the percentage of Satb2-positive neurons in the progeny of the targeted, EdU-labeled cNPCs, 4 days after electroporation of control construct (white column) or conditional Sox9 expression construct (red column) and 3 days after EdU administration at E14.5. **(O)** Quantification of the number of Satb2 and EdU double-positive, RFP-negative nuclei in the CP per microscopic field of 200-µm width, 4 days after electroporation of control construct (white column) or conditional Sox9 expression construct (red column) and 3 days after EdU administration at E14.5. **(G, H, L, M)** Scale bars, 50 µm. **(D, E, F, I, J, K, N, O)** Two-tailed, unpaired *t*-test: * p<0.05, ** p<0.01, n.s. not significant. Data are the mean of 3 **(E, F, I, J, K, N, O)** and 4 **(D)** embryos, each from a different litter; for each embryo, two microscopic fields, each of either 200-µm **(D, E, F, N, O)** or 150-µm **(I, J, K)** width were counted, and the values obtained were averaged. Error bars represent SEM.

In order to identify the genes that were affected by conditional Sox9 expression, we performed differential gene expression (DGE) analysis by comparing the expression levels of only protein-encoding genes in conditionally Sox9 expressing vs control High RFP cells. This yielded 749 downregulated and 870 upregulated protein-encoding genes upon Sox9 expression (q<0.01) (Fig. 5B, **Supp. Table 1**). To gain insight into the cell biological processes affected by conditional Sox9 expression, we examined the 870 genes upregulated in Sox9-expressing cells by gene ontology (GO) term enrichment analysis for biological process (Fig. 5C) and cellular component (Fig. 5D). This analysis revealed that the highest enrichment scores most frequently included genes related to the ECM (Fig. 5C and D). Similarly, KEGG pathway analysis using the same gene set as input yielded the highest enrichment scores for ECM production, organization and degradation (Fig. 5E).

In addition to the upregulation of expression of ECM components, conditional Sox9 expression induced the expression of gliogenic genes such as *Olig2* and *S100b* (Supp. Fig. 7A and B), corroborating the effect of conditional Sox9 expression on gliogenesis observed by immunohistochemistry (Fig. 4E and F).

**Fig. 7.**
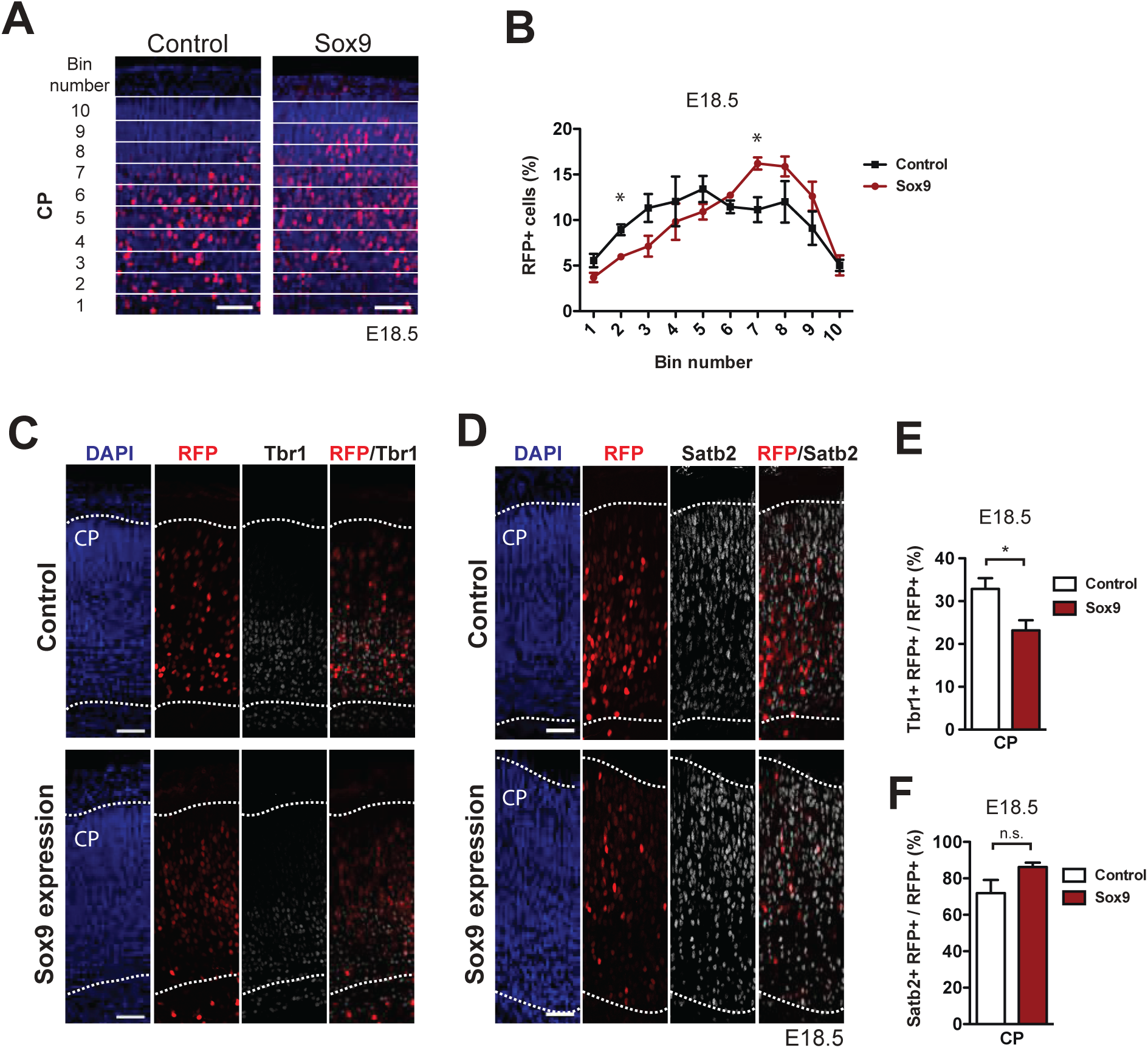
The neurons derived from mouse BPs targeted by conditional Sox9 expression are preferentially destined to the upper layers of the neocortex. Heterozygous *Tis21*-CreER^T2^ mouse embryos received tamoxifen at E12.5 and E13.5, were subjected to *in utero* electroporation of the neocortex at E13.5 with either control construct or conditional Sox9 expression construct, and subjected to immunostaining analyses at E18.5. **(A)** Immunofluorescence for RFP (red), combined with DAPI staining (blue), in the CP 5 days after electroporation of control or conditional Sox9 expression construct (see Fig. 3A). For the quantification of the distribution of RFP-positive nuclei (see **B**), images were divided into ten bins of equal size (numbered 1-10 from apical to basal) that span the entire CP. **(B)** Quantification of the percentage of the RFP-positive nuclei in the CP that are found in each bin (see **A**), 5 days after electroporation of control construct (black line) or conditional Sox9 expression construct (red line). **(C, D)** Double immunofluorescence for RFP (red) and either Tbr1 (white) **(C)** or Satb2 (white) **(D)**, combined with DAPI staining (blue), in the CP 5 days after electroporation of control or conditional Sox9 expression construct. Dotted lines delineate the apical and basal borders of the CP. **(E, F)** Quantification of the percentage of the RFP-positive nuclei in the CP that are Tbr1-positive **(E)** and Satb2-positive **(F)**, 5 days after electroporation of control construct (white columns) or conditional Sox9 expression construct (red columns). **(A, C, D)** Scale bars, 50 µm. **(B, E, F)** Two-tailed, unpaired *t*-test: * p<0.05, n.s. not significant. Data are the mean of 3 embryos, each from a different litter; for each embryo, two microscopic fields, each of either 200-µm **(B)** or 150-µm **(E, F)** width, were counted, and the values obtained were averaged. Error bars represent SEM.

### Conditional Sox9 expression induces cell non-autonomous BP proliferation resulting in increased neuron production, especially of upper-layer neurons

ECM components have been reported to increase the proliferative capacity of BPs, and cell non-autonomous effects have been considered in this context (Fietz et al., 2010; Fietz et al., 2012; Florio et al., 2015; Long et al., 2016; Long and Huttner, 2019; Stenzel et al., 2014). In light of this, we examined if upregulated ECM production had an effect on the progeny in the SVZ derived from non-electroporated cells. To this end, we analyzed the immunofluorescence of electroporated neocortex to compare control and conditional Sox9 expression with regard to (i) Ki67-positive cells in the SVZ that were RFP-negative, i.e. progeny derived from non-electroporated cells (Fig. 6A), (ii) RFP-negative abventricular mitoses (Fig. 6B), and (iii) RFP-negative Sox2-positive nuclei in the SVZ (Fig. 6C). Quantification per unit area of the numbers of Ki67-positive & RFP-negative nuclei in SVZ+IZ (Fig. 6D), of PH3-positive & RFP-negative abventricular mitoses (Fig. 6E), and of Sox2-positive & RFP-negative nuclei in SVZ+IZ (Fig. 6F) revealed significant increases for all three cell populations in conditionally Sox9-expressing neocortex. These findings indicate that conditional Sox9 expression in mouse BPs not only increases their proliferation cell-autonomously (see Fig. 3), but also results in a cell non-autonomous stimulation of BP proliferation.

We next investigated the outcome of the increased cell non-autonomous BP proliferation on neuron production. Specifically, we performed immunofluorescence for NeuN at E17.5 of mouse neocortex electroporated at E13.5 (Fig. 6G and H), and found that the percentage of NeuN-positive nuclei in the CP was significantly increased upon conditional Sox9 expression (Fig. 6I). Further examination of the NeuN-positive nuclei in the CP showed that conditional Sox9 expression in BPs resulted in a significant increase in the percentage of nuclei that were RFP-negative and NeuN-positive (Fig. 6K), whereas no significant change in the percentage of RFP-positive nuclei that were NeuN-positive was observed (Fig. 6J). Taken together, we conclude that conditional Sox9 expression in mouse BPs results in increased neuron production from BPs whose proliferation was stimulated in a cell non-autonomous manner.

We further dissected the effects of conditional Sox9 expression in mouse BPs with regard to the identity of the neurons produced (Lodato and Arlotta, 2015). To this end, we carried out immunofluorescence for the upper-layer marker Satb2 (Britanova et al., 2008; Leone et al., 2015) and the deep-layer marker Tbr1 (Hevner et al., 2001; Molyneaux et al., 2007) at E17.5 of mouse neocortex electroporated at E13.5 and subjected to EdU pulse-labeling at E14.5 (Fig. 6L and M, Supp Fig. 8A and B). On the one hand, conditional Sox9 expression in BPs resulted in a significant, more than two-fold increase in the number EdU-containing (EdU+) cells in the CP that were Satb2-positive and RFP-negative, i.e. progeny derived from of non-electroporated cells (Fig. 6O). On the other hand, no change in the proportion of EdU-containing (EdU+) RFP-positive (RFP+) cells that were Satb2-positive was observed upon conditional Sox9 expression in BPs (Fig. 6N). In contrast, the proportion of EdU-containing (EdU+) RFP-positive (RFP+) cells that were Tbr1-positive (Tbr1+) was found to be massively decreased upon conditional Sox9 expression in BPs (Supp Fig. 8C). These findings therefore not only indicate that upon conditional Sox9 expression in mouse BPs the production of deep-layer neurons is decreased and that of upper-layer neurons is increased, but also imply that the former decrease reflects an effect of Sox9 expression in the targeted BPs whereas the latter effect involves the non-targeted BPs whose proliferation is stimulated in a cell non-autonomous manner.

**Fig. 8.**
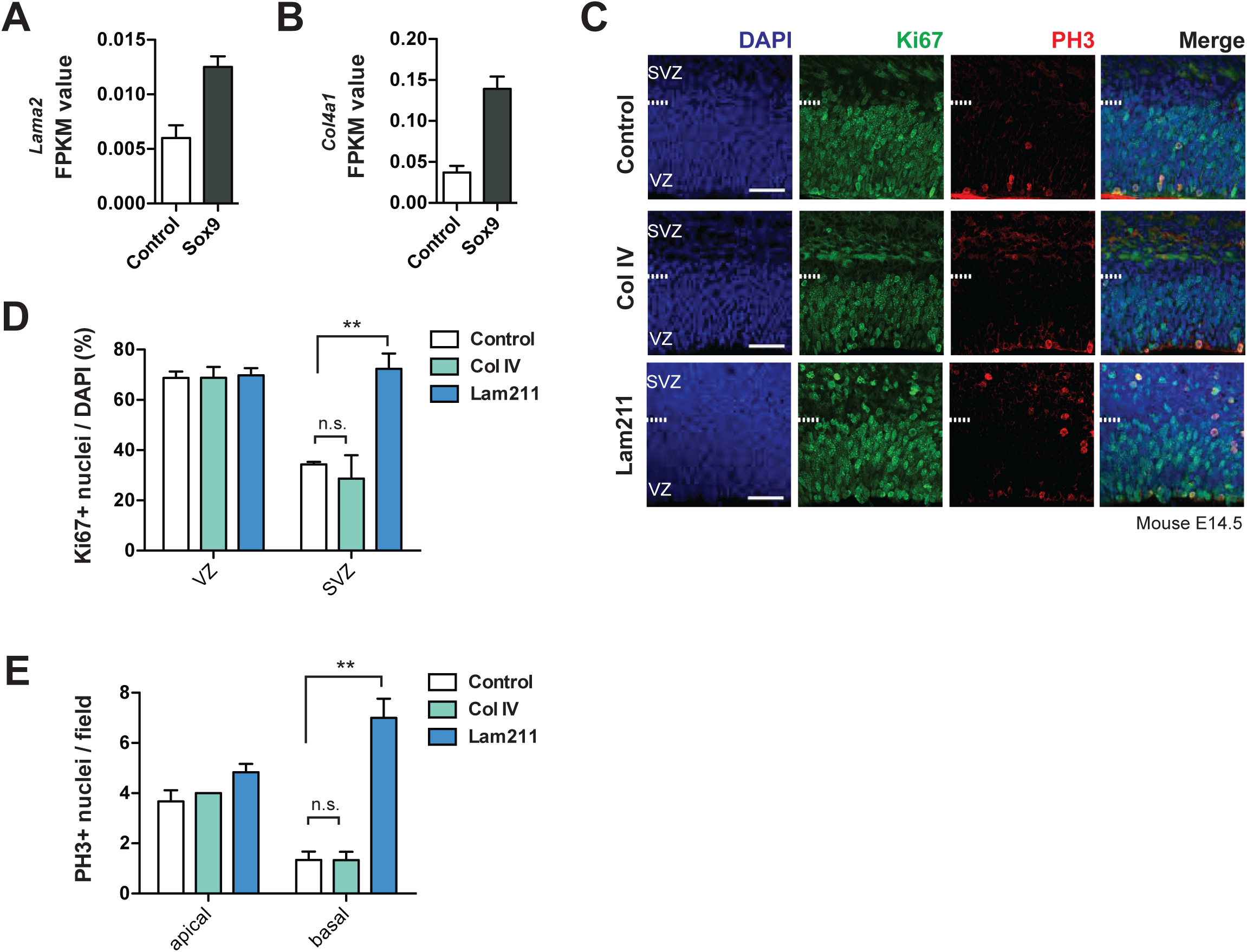
Laminin 211 induces BP proliferation in embryonic mouse neocortex. **(A, B)** FPKM values for *Lama2* mRNA (**A**) and *Col4a1* mRNA (**B**) in the high-level RFP-expressing cells at E15.5 upon electroporation of heterozygous *Tis21*-CreER^T2^ mouse embryos with control and conditional Sox9 expression constructs at E13.5 (for details, see Fig. 5A and legend). Data are the mean of transcriptomes of high-level RFP+ cells from 3 (control electroporated) and 4 (Sox9 electroporated) mouse embryonic neocortices, respectively. Error bars represent SEM. **(C)** Double immunofluorescence for Ki67 (green) and phosphohistone H3 (PH3, red), combined with DAPI staining (blue), of mouse E14.5 organotypic slices of neocortex cultured for 24 hours without (control) or with collagen IV (Col IV) or laminin α_2_β_1_ɣ_1_ (Lam211). Ventricular surface is down, dashed lines indicate the border between the VZ and SVZ. Scale bars, 50 µm. **(D)** Quantification of the percentage of the nuclei (identified by DAPI staining) that are Ki67-positive, in the VZ and SVZ of 24-hour control (white columns), Col IV-treated (green columns) and Lam211-treated (blue columns) E14.5 mouse neocortical organotypic slice cultures. **(E)** Quantification of the number of ventricular (apical) and abventricular (basal) PH3-positive mitoses, per microscopic field of 200-µm apical width, in 24-hour control (white columns), Col IV-treated (green columns) and Lam211-treated (blue columns) E14.5 mouse neocortical organotypic slice cultures. **(D, E)** Two-tailed, unpaired *t*-test, ** p<0.01. Data are the mean of 3 embryos, each from a different litter; for each embryo, two microscopic fields, each of 200-µm apical width, were counted, and the values obtained were averaged. Error bars represent SEM. Note that the three data for apical mitoses upon collagen IV treatment were identical.

We complemented these data by investigating the distribution of RFP-positive cells across the CP at E18.5, with the CP divided into 10 bins of equal size, comparing control and conditionally Sox9-expressing mouse neocortex electroporated at E13.5. (Fig. 7A). Quantification of the percentage of RFP+ cells in each bin showed that upon conditional Sox9 expression in BPs, the relative distribution of the RFP+ cells was shifted towards the upper layers of the CP, with a significant decrease in the percentage of RFP+ cells in the lower layers of the CP (specifically bin number 2, Fig. 7B) and an increase in the percentage of RFP+ cells in the upper layers of the CP (specifically bin number 7, Fig. 7B). Consistent with these data and with the observed decrease in the proportion of EdU-containing RFP-positive cells that were Tbr1-positive at E17.5 upon conditional Sox9 expression (Supp Fig. 8C), the proportion of all RFP+ cells in the CP that were Tbr1+ at E18.5 was significantly decreased upon conditional Sox9 expression (Fig. 7C and E). However, this decrease did not translate into a statistically significant increase in the proportion of RFP+ cells in the CP at E18.5 that were Satb2+ upon conditional Sox9 expression (Fig. 7D and F). Again, this was consistent with the lack of a statistically significant increase in the proportion of EdU-containing RFP-positive cells in the CP at E17.5 that were Satb2-positive upon conditional Sox9 expression in BPs (Fig. 6N), and with the notion that the increase in Satb2-positive neurons upon conditional Sox9 expression in BPs pertains mostly to progeny derived from of non-electroporated cells (Fig. 6O).

### ECM component laminin 211 promotes BP proliferation

Finally, in light of both, the increased expression of specific ECM components and the increase in BP proliferation, upon conditional Sox9 expression, we sought to directly demonstrate that selected candidate ECM components promote mouse BP proliferation. To this end, we incubated organotypic slices of E14.5 wildtype mouse neocortex in the absence and presence of either recombinant collagen IV or recombinant laminin 211 (laminin α2, β1, ɣ1). Collagen IV and laminin 211 were chosen as representative members of the collagen and laminin families of proteins, respectively, which have previously been implicated in cortical progenitor self-renewal and proliferation (Fietz et al., 2012). Of note, collagen IV mRNAs have been found to be expressed not only in the fetal human VZ but also ISVZ and OSVZ (Fietz et al., 2012), in line with the proliferative capacity of the cortical progenitors therein. Accordingly, collagen IV mRNAs are found to be expressed in both aRG and bRG (Florio et al., 2015). Likewise, with regard to laminin 211, laminin α2 (i) has been shown to be expressed not only in the embryonic mouse VZ but also SVZ (Lathia et al., 2007) and, notably, in the human OSVZ (Fietz et al., 2012), as well as in human aRG and bRG (Florio et al., 2015); and (ii) to be involved in the regulation of AP proliferation (Loulier et al., 2009).

The present analyses provided additional data that were at least consistent with a potential role of laminin 211 and collagen IV in promoting mouse BP proliferation upon conditional Sox9 expression. Specifically, in our analysis of the protein-encoding genes upregulated upon conditional Sox9 expression, the laminin α2 mRNA – although not among the list of the 870 upregulated genes – showed a ≈two-fold increase upon conditional Sox9 expression (Fig. 8A). Similarly, the collagen 4α1 mRNA, which is among the list of the 870 upregulated genes (**Supp. Table 1**), showed a >3-fold increase upon conditional Sox9 expression (Fig. 8B).

Following the treatment of organotypic slices of E14.5 wildtype mouse neocortex without or with recombinant collagen IV or laminin 211, our analysis of cell proliferation by immunofluorescence for Ki67 and phosphohistone H3 (Fig. 8C) revealed that laminin 211, but not collagen IV, caused a significant increase in the proportion of cycling cells (Ki67+) specifically in the SVZ, but not VZ (Fig. 8D). In addition, the number of basal mitoses (PH3+) per unit area was strikingly increased by laminin 211, but not by collagen IV, whereas the number of apical mitoses was largely unaffected by either ECM component (Fig. 8E). These findings provide strong support for the concept that conditional Sox9 expression in mouse BPs increases the expression of specific ECM components, notably laminins, which potentially promote BP proliferation in an autocrine (RFP-positive BPs, cell autonomous) and paracrine (RFP-negative BPs, cell non-autonomous) manner.

## Discussion

The present study establishes Sox9 as a master regulator of BP proliferation and cell fate. Remarkably, the four major effects of Sox9, that is, on (i) transcription, (ii) BP proliferation, (iii) neuron production, and (iv) BP cell fate, constitute a harmonic quartet of impacts, as is addressed one by one below.

### Sox9 is a major inducer of ECM components

Corroborating our considerations at the start of this study, i.e. that Sox9 – when expressed in the SVZ – could be a promising candidate to drive the expression of ECM components in BPs, our transcriptome analysis upon conditional Sox9 expression in mouse BPs revealed that, indeed, the 870 genes upregulated in Sox9-expressing BPs were predominantly related to the ECM by GO term enrichment analysis. The finding that Sox9 expression in BPs drives the expression of ECM components suggests an elegant explanation for the cell non-autonomous stimulation of BP proliferation by Sox9, as delineated below.

### Sox9 promotes BP proliferation cell-autonomously and non-autonomously

Comparison of embryonic mouse, embryonic ferret and fetal human neocortex showed that Sox9 expression in the SVZ correlated with the occurrence of BPs endowed with proliferative capacity, being detected in the ferret and human but not mouse SVZ. Further evidence in support of a positive role of Sox9 in the proliferation of ferret and human BPs was provided by the observations that these BPs were cycling and capable of cell cycle re-entry, and that they largely lacked expression of Tbr2, a marker that is often expressed in those BPs that are committed to neuron production rather than proliferation (Arnold et al., 2008; Englund et al., 2005; Florio et al., 2015; Kowalczyk et al., 2009; Sessa et al., 2008). Moreover, essentially all of the human bRG, the BP cell type particularly implicated in neocortical expansion (Fietz et al., 2010; Florio and Huttner, 2014; Hansen et al., 2010), were found to express Sox9. Taken together, all these findings were consistent with the notion that Sox9 promotes the proliferative capacity of BPs.

The conditional expression of Sox9 in mouse BPs, which normally lack proliferative capacity, provided direct evidence for this notion. Remarkably, however, the BPs that showed increased proliferative capacity upon conditional Sox9 expression were not only the progeny of electroporated cells, but also the progeny of the non-electroporated cells. In other words, Sox9 promoted not only the proliferation of those mouse BPs in which it was expressed in a targeted manner, but also of neighboring BPs lacking Sox9 expression. Previous findings have implicated ECM components as stimulators of cNPC proliferation that act via integrin signaling (Arai et al., 2011; Fietz et al., 2010; Fietz et al., 2012; Florio et al., 2015; Long et al., 2016; Stenzel et al., 2014). Moreover, the present observations demonstrate that Sox9 expression in BPs induces the expression of ECM components, some of which (like laminin 211) can promote BP proliferation. It therefore seems likely that the cell non-autonomous stimulation of mouse BP proliferation upon conditional Sox9 expression is due to ECM components that have been secreted from Sox9-expressing BPs and then act on neighboring BPs that themselves have not been targeted to express Sox9. If so, this interplay reflects how well the Sox9 effects on transcription and BP proliferation are orchestrated.

We do not know which genes whose expression is increased upon conditional Sox9 expression in mouse BPs are responsible for the cell-autonomous stimulation of BP proliferation by Sox9. However, given the cell non-autonomous stimulation of BP proliferation upon Sox9 expression which likely involves ECM components secreted from Sox9-expressing BPs, it seems possible that the mechanism underlying the cell-autonomous stimulation of BP proliferation by Sox9 also involves ECM components secreted from the Sox9-expressing BPs and acting on them in an autocrine fashion.

### Sox9 increases production of upper-layer neurons

The effects of conditional Sox9 expression in mouse BPs on neuron production added yet a third line of evidence for the orchestrated effects of Sox9. An increase in upper-layer neuron production is considered to be one of the hallmarks of neocortical expansion (Hutsler et al., 2005; Molnar et al., 2006). Accordingly, conditional Sox9 expression in mouse BPs resulted in this phenotype. Interestingly, however, whereas the Sox9-induced decrease in the production of deep-layer neurons was due to the effects of Sox9 in the Sox9-expressing BPs, the increase in the production of upper-layer neurons reflected an indirect effect via the BPs that themselves were not targeted to express Sox9. This illustrates how a synergistic effect (less deep-layer neurons, more upper-layer neurons) can be achieved via concerted direct and indirect effects of Sox9 on BPs.

### Sox9 switches BPs to gliogenesis

Finally, the quartet of harmonic impacts of Sox9 expression was completed by its effects on the fate of BPs. Specifically, not only did conditional Sox9 expression in mouse BPs reduce the proportion of Tbr2-positive BPs, consistent with the stimulation of BP proliferation by Sox9, but conditional Sox9 expression also induced Olig2 in a subset of BPs, indicative of switching them to gliogenesis. In the adult mouse brain, Sox9 has been reported to be expressed simultaneously by mature astrocytes and neurogenic SVZ progenitors (Cheng et al., 2009; Martini et al., 2013; Nagao et al., 2014; Nait-Oumesmar et al., 2008; Sun et al., 2017). In the embryonic mouse neocortex, Sox9-expressing aRG have been shown to be capable of generating neurons for all layers (Kaplan et al., 2017). Our findings are therefore consistent with the multi-faceted role of Sox9 in neurogenesis and gliogenesis (Cheng et al., 2009; Kaplan et al., 2017; Nagao et al., 2014; Nagao et al., 2016; Selvaraj et al., 2017; Sun et al., 2017). As neocortical expansion is characterized not only by an increase in neurogenesis, but also in gliogenesis (Rash et al., 2019), these effects of Sox9 expression again highlight the concerted nature of its action. Hence, when collectively considering the effects of Sox9 on transcription, BP proliferation and fate, and neuron production, this transcription factor shows all the hallmarks expected for a master regulator that acts in the SVZ to promote neocortical expansion.

## Materials and Methods

### Neocortical tissue

Embryos from C57Bl/6JOlaHsd mice at embryonic day (E) 13.5 and 14.5 were used as wild type. Conditional Sox9 expression experiments were carried out in heterozygous embryos of the *Tis21*-CreER^T2^ transgenic mouse line, in which exon 1 of the *Tis21* gene was replaced with a CreER^T2^ cassette (Wong et al., 2015). E0.5 was specified as the noon of the day when plugged females were observed. All mouse lines used in this study were congenic and kept in pathogen-free conditions at the Biomedical Services facility of the Max Planck Institute of Molecular Cell Biology and Genetics, Dresden. All experiments were approved by the Governmental Landesdirektion Sachsen. License numbers for the experiments performed are: 24–9168.11-9/2009-2 (for in utero electroporation, tamoxifen and EdU administration) and 24–9168.24-9/2012-1 (for tissue collection).

Pregnant ferrets were obtained from Marshall BioResources and housed at the BioCrea GmBH (Radebeul, Saxony, Germany). Several days prior to embryo collection, ferrets were transferred to the Biomedical Services facility of the Max Planck Institute of Molecular Cell Biology and Genetics, Dresden. All animal experiments are conducted according to the German animal welfare legislation.

Human fetal neocortical tissue (specifically for Fig. 1C and I) was provided by the Klinik und Poliklinik für Frauenheilkunde und Geburtshilfe, Universitätsklinikum Carl Gustav Carus, involving elective pregnancy terminations with informed written maternal consents and approval by the local University Hospital Ethical Committees. Human fetal neocortical tissue (specifically for Fig. 2F) was also obtained from Novogenix Laboratories (Torrance, CA). The age of fetuses was assessed by ultrasound measurements of crown-rump length and other standard criteria of developmental stage determination.

### Plasmids

Plasmid pCAGGS-LoxP-membraneGAP43-GFP-LoxP-IRES-nRFP (referred as control construct) was generated as previously described (Wong et al., 2015) and kindly provided by Dr. Fong Kuan Wong. Plasmid pCAGGS-LoxP-membraneGAP43-GFP-LoxP-Sox9-IRES-nRFP (referred as conditional Sox9 expression construct) was generated by cloning the Sox9 coding sequence into the control construct 5’ to the IRES sequence. The Sox9 coding sequence was amplified from FUW-TetO-Sox9 plasmid (Addgene, #41080) and cloned into the control plasmid using restriction enzyme XhoI (NEB, #R146L). pCAGGS-Cre plasmid was generated as described (Kranz et al., 2010) and kindly provided by Dr. JiFeng Fei.

Primers used to amplify the Sox9 coding sequence:

Sox9_F: 5’-TCTCGAGGCCGCCATGAATCTCCTGGACCCCTTC-3’

Sox9_R: 5’-TCTCGAGTCAGGGTCTGGTGAGCTGTGT-3’

### Cell culture

HEK293T cells were plated in 24-well plates with a density of 5×10^4^ cells per well, and cultured in DMEM containing 10% fetal bovine serum. After 24 hours of culturing, the cells were transfected using Lipofectamine 2000 (Invitrogen) with 1 μg of conditional Sox9 expression plasmid, with 1 μg of control plasmid, or with either conditional Sox9 expression or control plasmid in combination with 1 μg of pCAGGS-Cre plasmid. After transfection, the cells were cultured for an additional 48 hours and fixed with 4% (wt/vol) paraformaldehyde in 120 mM phosphate buffer (pH 7.4) (referred to from here on simply as PFA) for 10 min at room temperature.

### *In utero* electroporation

*In utero* electroporations (IUEPs) were performed as previously described (Shimogori and Ogawa, 2008). Pregnant *Tis21*-CreER^T2^ mice were treated via oral gavage with 2 mg tamoxifen dissolved in corn oil, at E12.5 and at E13.5. After tamoxifen administration, pregnant mice carrying E13.5 embryos were anesthetized with isofluorane. Then, 1.5 μg/μl of either control or conditional Sox9 expression plasmid DNA was mixed with 0.25% Fast Green FCF dye and was intraventricularly injected into each embryo with a glass microcapillary and electroporated by a series of 6 pulses (30V) with 1 ms intervals. Embryos were collected at 24 h, 48 h, 4 days or 5 days after electroporation.

### EdU labeling

EdU was dissolved in PBS at a concentration of 1 mg/ml. For analyzing the cell cycle re-entry of Sox9-expressing progenitors, P0 ferret kits were injected intraperitoneally with 100 μl of EdU solution. P2 ferret kits were subjected to hypothermic anesthesia in crushed ice and sacrificed by intracardiac perfusion with 4% PFA at 37°C. In order to analyze the cell cycle re-entry of the progeny of electroporated progenitors in mouse embryos, 100 μl of EdU solution was intraperitoneally injected into pregnant, electroporated mice at E14.5. EdU fluorescence was detected using the Click-iT EdU Alexa Fluor 647 Imaging Kit (Invitrogen C10340) according to the manufacturer’s instructions.

### Immunofluorescence

For immunofluorescence of transfected HEK293T cells, the cells were fixed on coverslips with 4% PFA at room temperature for 10 min and permeabilized with 0.3% Triton X-100 in PBS for 30 min, then quenched with 0.1 M glycine in 1x PBS for 30 min. Coverslips were then washed with 1x PBS containing 0.2% gelatin, an additional 300 mM NaCl, and 0.3% Triton X-100 and incubated with primary antibodies either for 3 hours at room temperature or overnight at 4°C, followed by incubation with secondary antibodies at room temperature for 1 hour. Cells were washed with PBS and mounted in Mowiol (Merck Biosciences).

In order to perform immunofluorescence on neocortex, the tissue was fixed with 4% PFA overnight at 4℃. The tissue was further processed for either vibratome or cyrosectioning. Vibratome sections were obtained from neocortical tissue embedded in 3% low melting agarose using a Leica 1000 vibratome. The sections were 50 μm thick and stored in PBS for further processing. For cryosectioning, neocortical tissue was incubated in a 30% sucrose solution overnight at 4°C, embedded in Tissue-TEK OCT compound (Sakura Finetek) and stored at −20°C. Cryosections were 10-16 μm thick.

The sections were subjected to antigen retrieval by incubation in 0.01 M sodium citrate (pH 6.0) in a water bath at 70°C for 1 hour. Sections were then incubated with 0.3% Triton X-100 in PBS for 30 min for permeabilization, followed by 0.1 M glycine for 30 min for quenching. Afterwards, sections were incubated overnight at 4°C with primary antibodies in 1x PBS containing 0.2% gelatin, an additional 300 mM NaCl and 0.3% Triton X-100. The sections were incubated with secondary antibodies for 1 hour at room temperature. The following primary antibodies were used in this study: rabbit anti-Sox9 (Sigma, HPA001758, 1:300), rat anti-RFP (ChromoTek, 5F8, 1:500), chicken anti-GFP (Aves labs, GFP-1020, 1:500), rabbit anti-Ki67 (Abcam, ab15580, 1:200), mouse anti-PCNA (Millipore, CBL407, 1:300), goat anti-Sox2 (Santa Cruz, sc-17320, 1:200), rabbit anti-Tbr2 (Abcam, ab23345, 1:200), sheep anti-Tbr2 (R+D Systems, AF6166, 1:500), mouse anti-Olig2 (Millipore, MABN50, 1:200), mouse anti-NeuN (Millipore, MAB377, 1:200), rat anti-phosphohistone H3 (Abcam, ab10543, 1:500), mouse anti-phosphovimentin (Abcam, ab22651, 1:300), mouse anti-Satb2 (Abcam, ab51502, 1:300), rabbit anti-Tbr1 (Abcam, ab31940, 1:200). The secondary antibodies used in this study were donkey- or goat-derived and were coupled to Alexa Fluor 405, 488, 555 and 647 (Life Technologies). Secondary antibody stainings (except when Alexa Fluor 405 was used) were combined with DAPI (Sigma).

### Image acquisition

Images were acquired using a Zeiss LSM700 confocal microscope with 20x, 40x, and 63x objectives, with pixel resolution of 1024×1024 or 2048×2048 and in stacks of 1.5 μm-thick optical sections. Images scanned in tiles were stitched using ZEN2012 software. All images were visualized and processed using Fiji software (https://fiji.sc/).

### Quantification and statistical analysis

All cell counts were performed using the Fiji software in standardized microscopic fields and calculated in Microsoft Office Excel. Statistics analyses were conducted using Prism (GraphPad Software) and unpaired Student’s t-test was used. Sample sizes, the statistical test and significance are indicated in each Figure legend.

### FACS

Dorsolateral cortices were dissected from hemispheres electroporated with control or conditional Sox9 expression plasmids. In each experiment, cortices from 1-3 embryos from 2-5 mothers were pooled and dissociated into single cell suspension using the MACS Neural Tissue Dissociation kit with papain (Miltenyi Biotec) according to manufacturer’s instructions. Dissociated cells were transferred into 5 ml tubes (Falcon) through a 35 μm cell-strainer cap. FACS was performed with a 5-laser-BD FACSAria Fusion (BD Bioscience) and analyzed with FACS Diva software v8.0 (BD Bioscience). First, live cells were identified based on their size and shape and a gate for live cells was set on the SSC/FSC dot-plot. Then a gate was set on FSC-W/FSC-H for sorting. Cells from a non-electroporated brain were used as negative control and fluorescence for RFP+ electroporated cells was detected. Out of the live cells (see above), single dot-plots were created for FSC-H/PE-Texas Red-A (yellow/green laser, 561 nm) and two gates were created for low intensity (low RFP) and high intensity RFP (high RFP) fluorescence. For qPCR experiments, 12,000 cells each from RFP–, low RFP and high RFP populations were sorted into 350 μl of RLT lysis buffer (QIAGEN) with 2 μl of β-mercaptoethanol. For RNA-sequencing experiments, 5,000 cells each from low RFP and high RFP populations were sorted into 350 μl of RLT lysis buffer with 2 μl of β-mercaptoethanol.

### qPCR

Total RNA from sorted cells was extracted using the QIAGEN RNeasy Micro kit according to the manufacturer’s protocol. cDNA was synthesized with random hexamers and Superscript III Reverse Transcriptase (Life Technologies, #18080044). qPCR experiments were performed on Mx3000P system (Agilent Technologies) using Absolute qPCR SYBR Green mix (Thermo Scientific). Relative mRNA levels were calculated using the comparative 2^-ΔΔCT^ method (Livak and Schmittgen, 2001). The housekeeping gene *Gapdh* was used as reference gene. Each experiment was performed in triplicate and with 4-5 biological replicates.

Primers used for qPCR experiments:

Sox9-F: 5’-AGGAAGCTGGCAGACCAGT-3’

Sox9-R: 5’-CTCCTCCACGAAGGGTCTCT-3’

Olig2-F: 5’-CCCCAGGGATGATCTAAGC-3’

Olig2-R: 5’-CAGAGCCAGGTTCTCCTCC-3’

Tbr2-F: 5’-GACCTCCAGGGACAATCTGA-3’

Tbr2-R: 5’-GTGACGGCCTACCAAAACAC-3’

Gapdh-F: 5’-TGAAGCAGGCATCTGAGGG-3’

Gapdh-R: 5’-CGAAGGTGGAAGAGTGGGAG-3’

### RNA sequencing

Total RNA from sorted cells was extracted using the QIAGEN RNeasy Micro kit according to the manufacturer’s protocol. cDNA synthesis was performed with SmartScribe reverse transcriptase (Clontech), universal poly-dT primers and template switching oligos. Purified cDNA was amplified with Advantage 2 DNA Polymerase in 12 cycles, and subjected to ultrasonic shearing with Covaris S2. Afterwards, standard Illumina fragment libraries were prepared using NEBnext chemistries (New England Biolabs). Library preparation consisted of fragment end-repair, A-tailing and ligation to indexed Illumina TruSeq adapters, followed by a universally primed PCR amplification of 15 cycles. Then, libraries were purified using XP beads (Beckman Coulter) and quantified by qPCR. Samples were subjected to Illumina 75-bp single end sequencing on an Illumina NextSeq platform.

### Transcriptome analysis

Reads of the same sample on different sequencing lanes were combined and processed by adapter trimming using cutadapt. Alignment of the processed reads to a mouse reference genome (mm10) was performed by STAR. RNA sequencing data was expressed as FPKM (fragments per kilobase of exon per million fragments mapped) values by quantification of the genes from Ensemble release 70 (**Supp. Table 2**). Replicates were clustered using Jensen-Shannon divergence and differential expression analysis was performed using DESeq2. Differential expression analysis was implemented with raw counts as input and differentially expressed genes were identified using a cutoff of q<0.01. Functional annotation clustering of the differentially expressed genes was performed using DAVID (https://david.ncifcrf.gov/) or Enrichr (http://amp.pharm.mssm.edu/Enrichr/) with default settings and using genes as input. Data visualization was performed using Cummerbund and R (http://www.r-project.org).

### Organotypic slice culture

Wildtype E14.5 mouse brains were first embedded in 3% low melting point agarose and 250 μm thick sections were obtained using a vibratome. The agarose on the sections was removed and slices were embedded in type Ia collagen (Cellmatrix, Nitta Gelatin) at a concentration of 1.5 mg/ml in DMEM and neutralizing buffer at room temperature, as described in the manufacturer’s protocol. Slices in the collagen mix were placed in 35 mm Petri dishes with a 14 mm microwell (MatTek Cooperation). Within a section the two hemispheres were separated, one hemisphere was embedded into type Ia collagen only as control, and the other was embedded into type Ia collagen mixed with either 0.1 mg/ml laminin 211 (Biolamina, human recombinant laminin LN211) or 0.1 mg/ml collagen IV (Abcam, Natural Human Collagen IV protein (FAM) ab123531). Petri dishes with embedded slices were kept at 37°C for 40 min to allow the collagen mix to set completely. Afterwards 2 ml of slice culture medium was added into each dish and the slices were cultured for 24 hours in a humidified incubation chamber supplied with 40% O_2_, 5% CO_2_ and 55% N_2_.

## Acknowledgments

We thank the Services and Facilities of the Max Planck Institute of Molecular Cell Biology and Genetics for their outstanding technical support, notably Jussi Helppi and his team of the Biomedical Services (BMS), Jan Peychl and his team of the Light Microscopy Facility and Lena Hersemann from the team of Scientific Computing Facility. We would like to thank Andreas Dahl and the Deep Sequencing Group of the DFG Research Center for Regenerative Therapies for performing the RNA-sequencing. We would like to thank, in particular, Fong Kuan Wong for kindly providing the control plasmid, Miguel Turrero-Garcia for help with perfusion of ferret kits, Alex Sykes for assistance on molecular cloning, Mareike Albert for help with FACS and RT-PCR, Milos Kostic and Nereo Kalebic for advice, Takashi Namba for his constructive comments on the manuscript, and all members of the Huttner group for helpful discussions. A.G. and M.F. were members of the International Max Planck Research School for Cell, Developmental and Systems Biology and doctoral students at the Technische Universität Dresden.

## Author contributions

AG: Conceptualization, Formal analysis, Investigation, Visualization, Methodology, Writing—original draft, Writing—review and editing, Performed *in utero* electroporations, cell sorting, RNA isolation, RT-PCR and gene expression analyses; DS: Conceptualization, Investigation, Methodology, Provided supervision to Ayse Güven; KRL: Methodology, Assisted organotypic slice cultures, Writing – input for original draft; MF: Methodology, Analysis and interpretation of the RNA-sequencing data and GO term analyses, Assisted human tissue dissection, Assisted FACS; HB: Analysis of the RNA-sequencing data; WBH: Conceptualization, Interpretation of data, Supervision, Funding acquisition, Project administration, Writing—original draft, Writing—review and editing.

## Supplementary Figure Legends

**Supp. Fig. 1.**
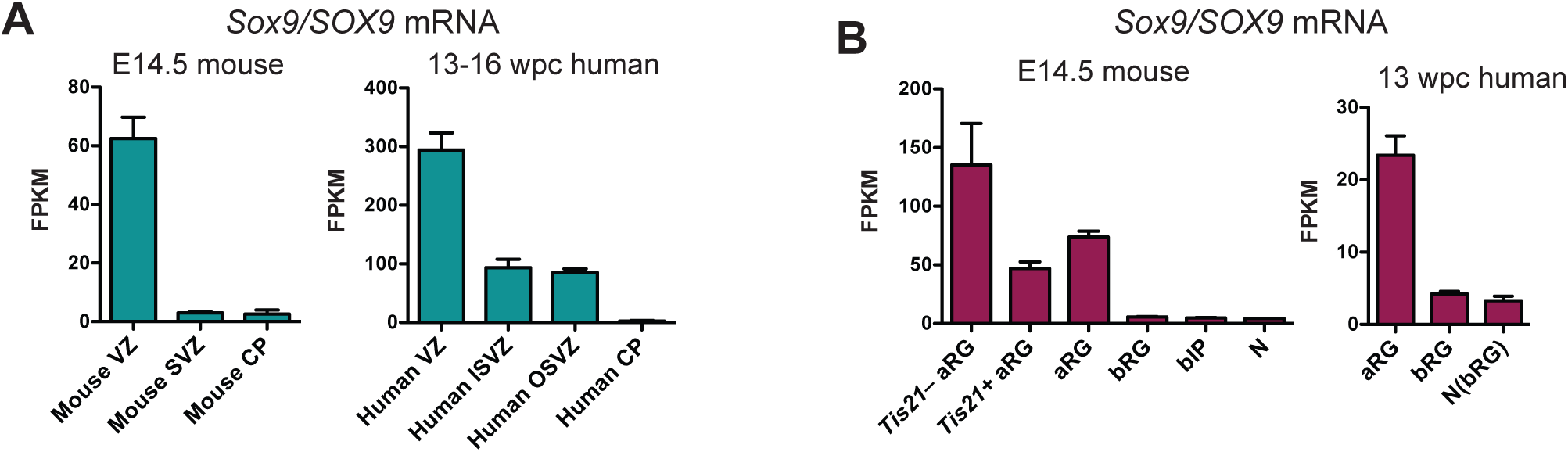
*Sox9* mRNA is expressed in fetal human, but not embryonic mouse, BPs. **(A)** FPKM values reflecting the *Sox9/SOX9* mRNA levels in the germinal zones of mouse E14.5 (left) and human 13-16 wpc (right) neocortex, as determined previously (Fietz et al., 2012). **(B)** FPKM values reflecting the *Sox9/SOX9* mRNA levels in distinct mouse E14.5 (left) and human 13 wpc (right) neocortical cell subpopulation, as determined previously (Florio et al., 2015). *Tis21* –/+, *Tis21*-GFP–negative/positive; bIP, mouse cell subpopulation containing bIPs and a minor proportion of other cell types that do not express prominin-1 and Tubb3 and are not labeled by DiI (e.g. endothelial cells); N, mouse neurons; N(bRG), human neuron fraction containing bRG in G1. Data are the mean of 5 (**A**) and 4 (**B**) mouse transcriptomes and of 6 **(A**, one 13 wpc, two 14 wpc, one 15 wpc and 2 16 wpc determination(s)) and 4 (**B**) human transcriptomes. Error bars represent SD.

**Supp. Fig. 2.**
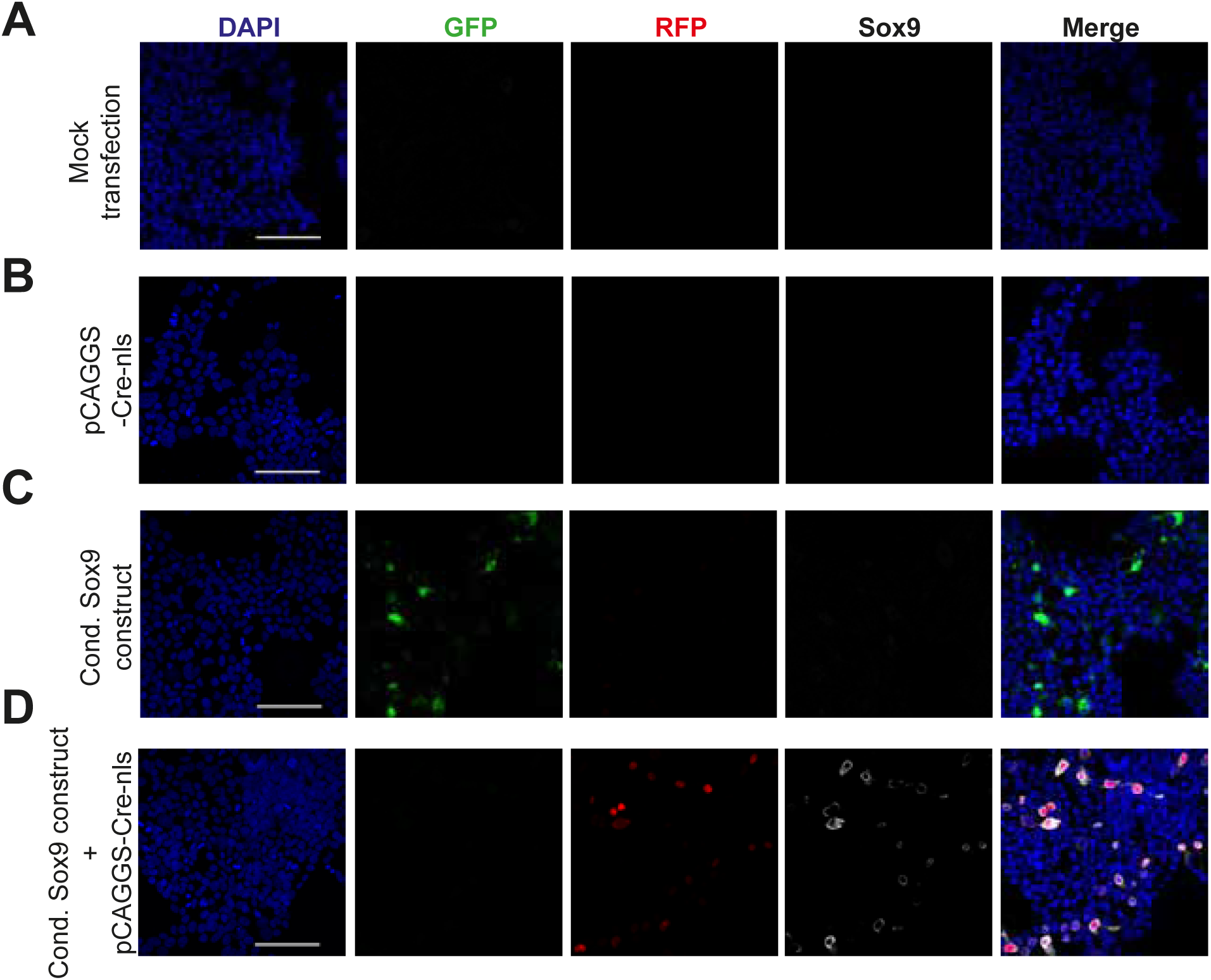
*In vitro* validation of the functionality of the conditional Sox9 expression construct using HEK293T cells. Triple immunofluorescence for GFP (green), RFP (red) and Sox9 (white), combined with DAPI staining (blue), of HEK293T cells subjected to mock transfection **(A)** or transfected with pCAGGS-Cre-nls **(B)**, the conditional (cond.) Sox9 expression construct **(C)**, or the conditional (cond.) Sox9 expression construct plus pCAGGS-Cre-nls **(D)**, followed by 48 hours of culture. Scale bars, 100 µm.

**Supp. Fig. 3.**
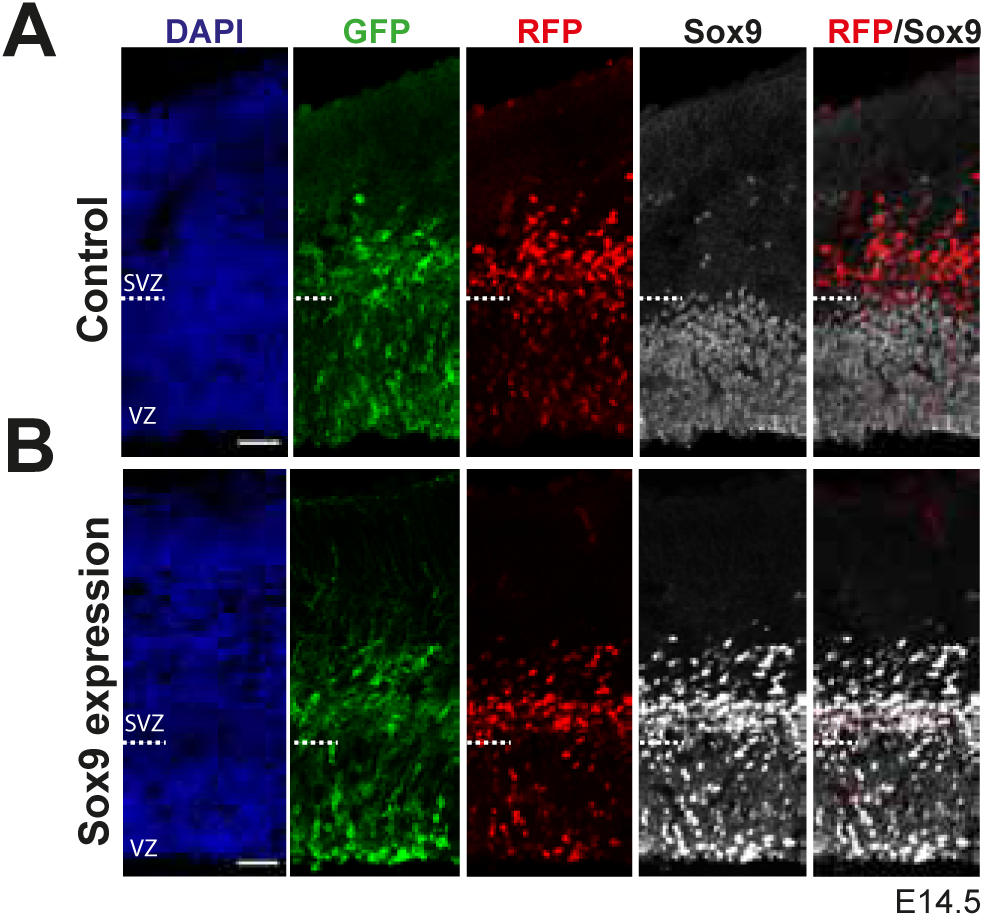
Validation of the Sox9 expression, elicited specifically in BP-genic aRGs and BPs upon *in utero* electroporation of the conditional Sox9 expression construct into the neocortex of *Tis21*-CreER^T2^ mouse embryos. Heterozygous *Tis21*-CreER^T2^ mouse embryos received tamoxifen at E12.5 and E13.5, and the neocortex was *in utero* electroporated with either control construct (**A**) or conditional Sox9 expression construct (**B**) (see Fig. 3A) at E13.5, followed by triple immunofluorescence for GFP (green), RFP (red) and Sox9 (white), combined with DAPI staining (blue), 24 hours later. Dashed lines indicate the border between VZ and SVZ. Scale bar, 50 µm.

**Supp. Fig. 4.**
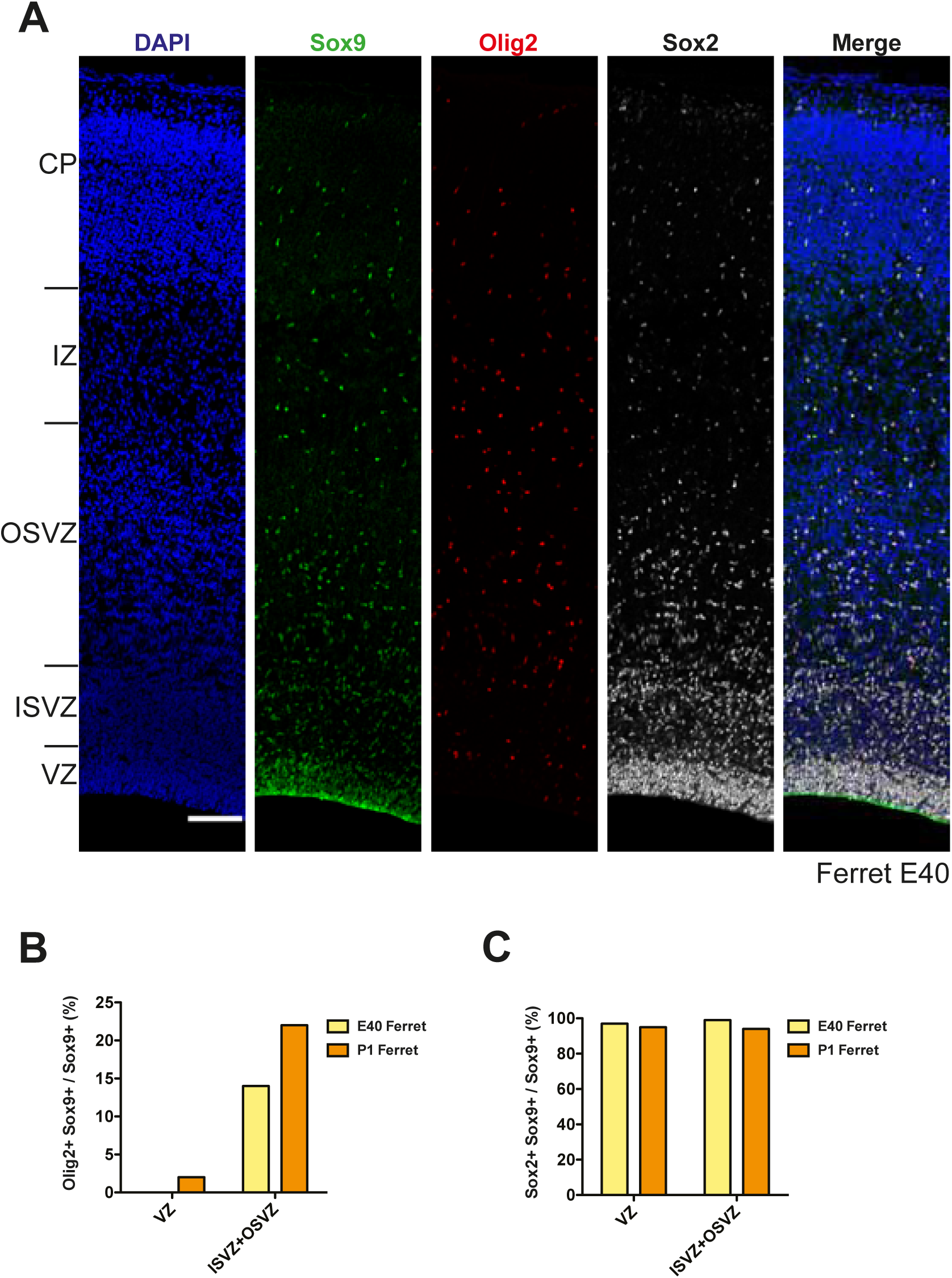
Gliogenic potential of Sox9-expressing cNPCs in the germinal zones of developing ferret neocortex. **(A)** Double immunofluorescence for Sox9 (green) and Olig2 (red), combined with DAPI staining (blue), of E40 ferret neocortex. Ventricular surface is down, upper margins of images correspond to the pial surface. Scale bar, 100 µm. **(B)** Quantification of the percentage of the Sox9-positive nuclei in the VZ, and of the Sox9-positive nuclei in the ISVZ plus OSVZ, that are Olig2-positive, in the E40 (yellow columns, quantified from the images shown in (**A**) and P1 (brown columns, images used for quantification are not shown) ferret neocortex.

**Supp. Fig. 5.**
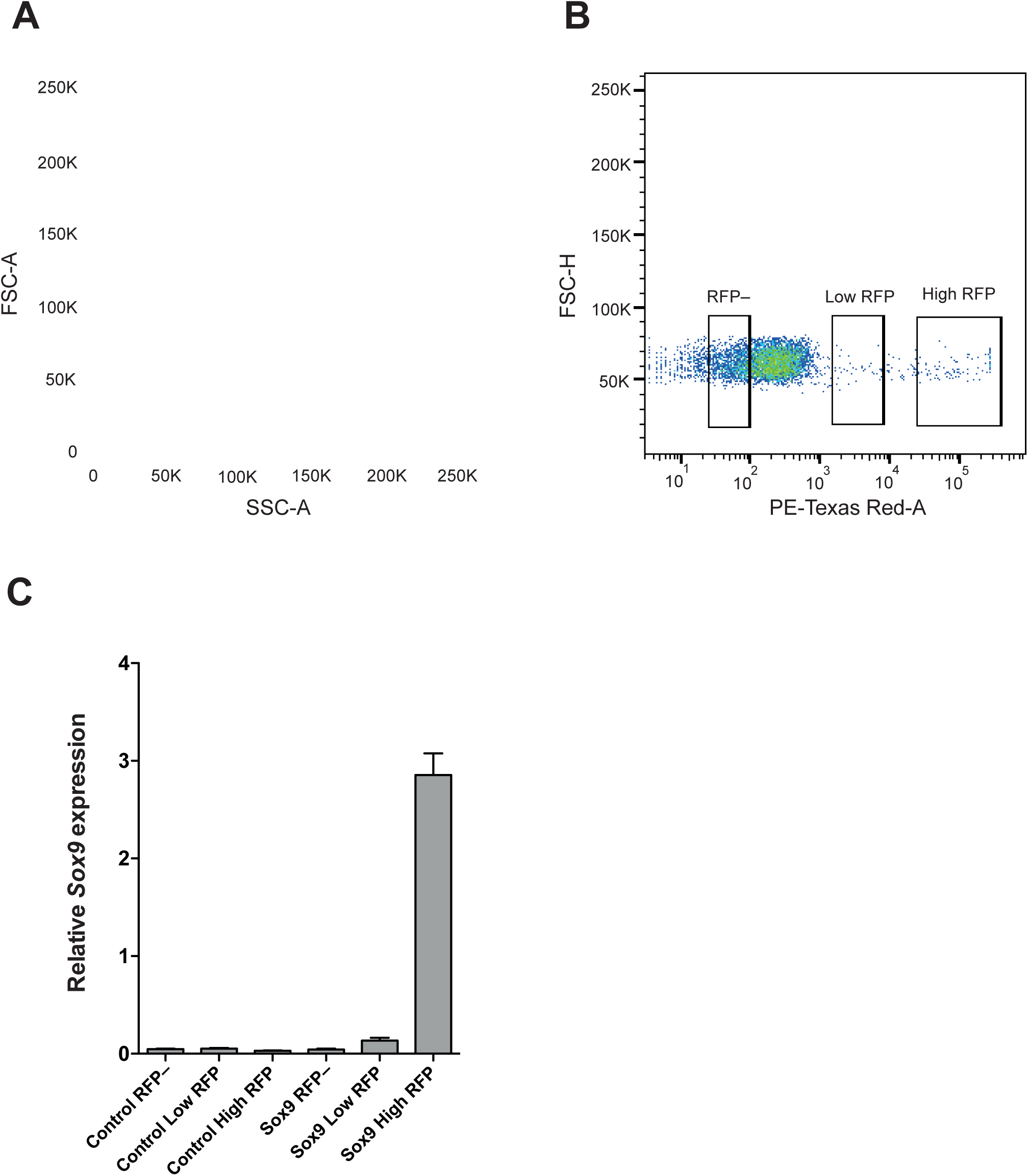
Isolation of RFP-negative, low-level RFP-expressing and high-level RFP-expressing cNPC subpopulations by FACS and determination of *Sox9* mRNA levels in the six subpopulations. **(A)** FSC/SSC dot-plot showing the distribution of dissociated cells, obtained from neocortices electroporated with control construct or conditional Sox9 expression construct (see Fig. 4H), based on their size and shape. A gate for live cells was set as indicated by the polygon. **(B)** FSC-H/PE-Texas Red-A dot-plot of the live cells (see **A**) showing the distribution of cells based on their RFP fluorescence intensity. Three gates were set in order to sort RFP-negative (left rectangle), low-level RFP-expressing (middle rectangle) and high-level RFP-expressing (right rectangle) subpopulations. **(C)** Quantification of *Sox9* mRNA levels relative to the *Gapdh* mRNA level by qPCR analysis in RFP-negative, low-level RFP-expressing and high-level RFP-expressing cell subpopulations isolated by FACS (see **A, B**) 48 hours after electroporation of control construct (left three columns) or conditional Sox9 expression construct (right three columns) (see Fig. 4H).

**Supp. Fig. 6.**
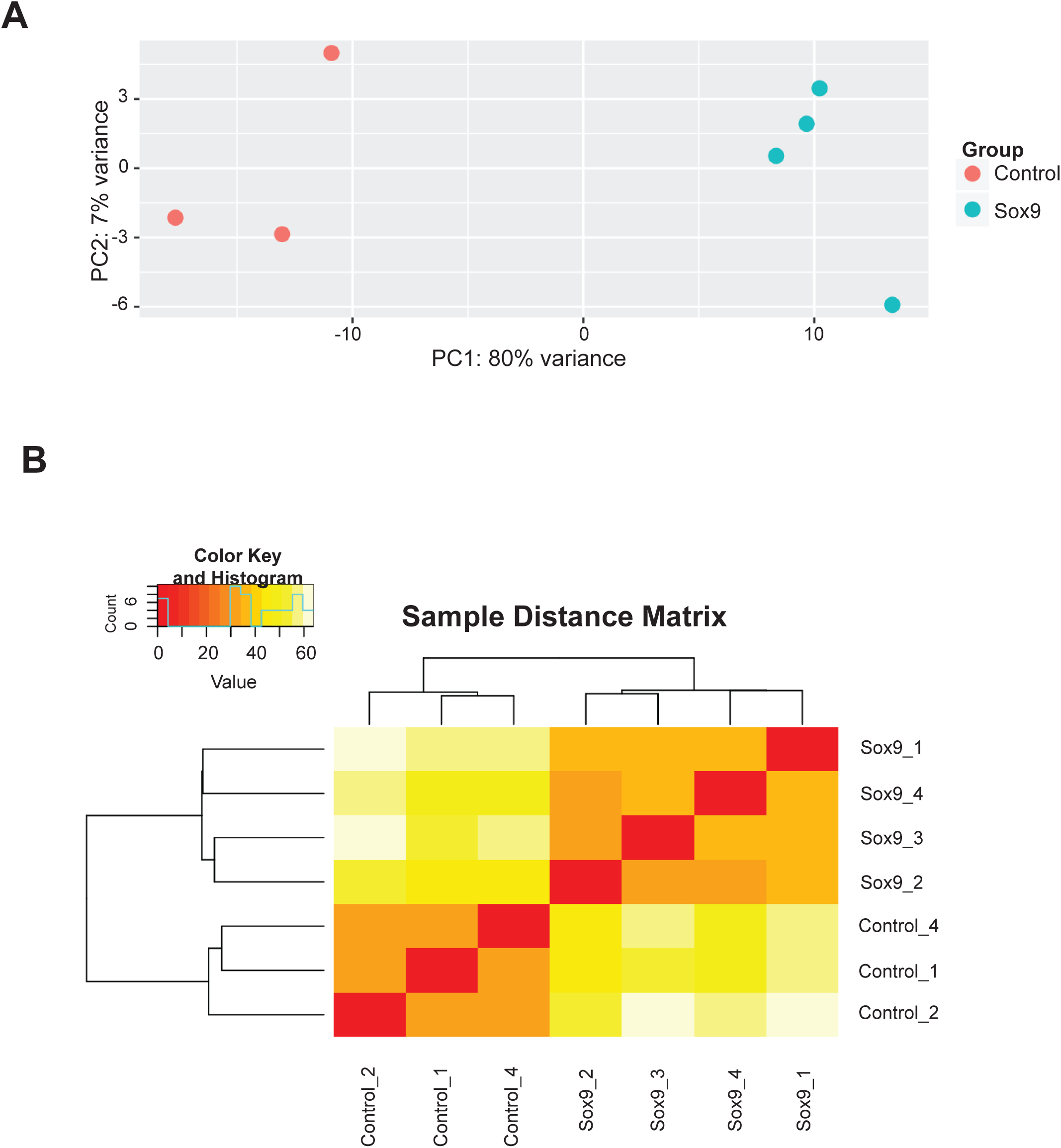
Principal component analysis and hierarchical clustering of electroporated cNPCs. **(A)** Principal component (PC) analysis of the high-level RFP-expressing cell subpopulations upon electroporation with control construct (3 replicates, red) and conditional Sox9 expression construct (4 replicates, turquoise) (see Fig. 5A). **(B)** Hierarchical clustering of the high-level RFP-expressing cell subpopulations upon electroporation with control construct (3 replicates) and conditional Sox9 expression construct (4 replicates) (see Fig. 5A).

**Supp. Fig. 7.**
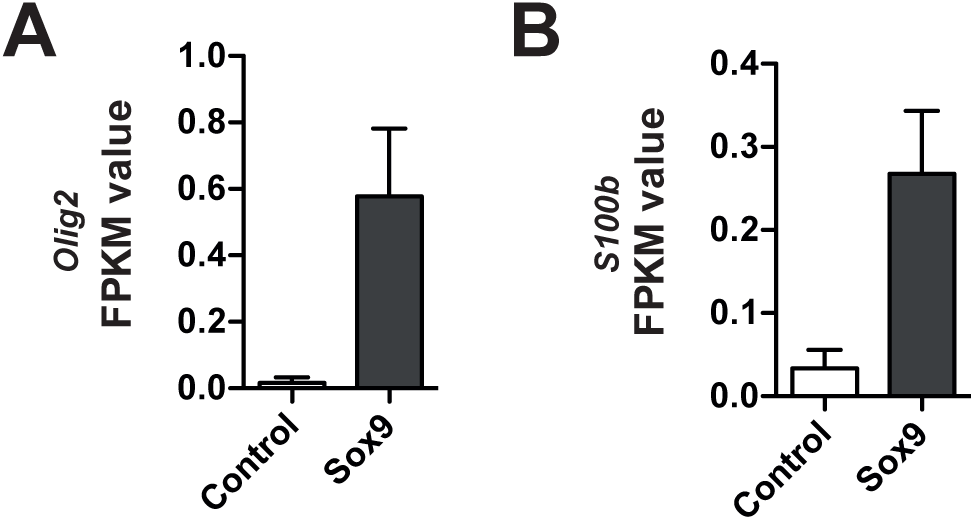
mRNAs of gliogenesis markers are upregulated upon conditional Sox9 expression. FPKM values for *Olig2* mRNA (**A**) and *S100b* mRNA (**B**) in the high-level RFP-expressing cells at E15.5 upon electroporation of heterozygous *Tis21*-CreER^T2^ mouse embryos with control and conditional Sox9 expression constructs at E13.5 (for details, see Fig. 5A and legend). Data are the mean of transcriptomes of high-level RFP+ cells from 3 (control electroporated) and 4 (Sox9 electroporated) mouse embryonic neocortices, respectively. Error bars represent SEM.

**Supp. Fig. 8.**
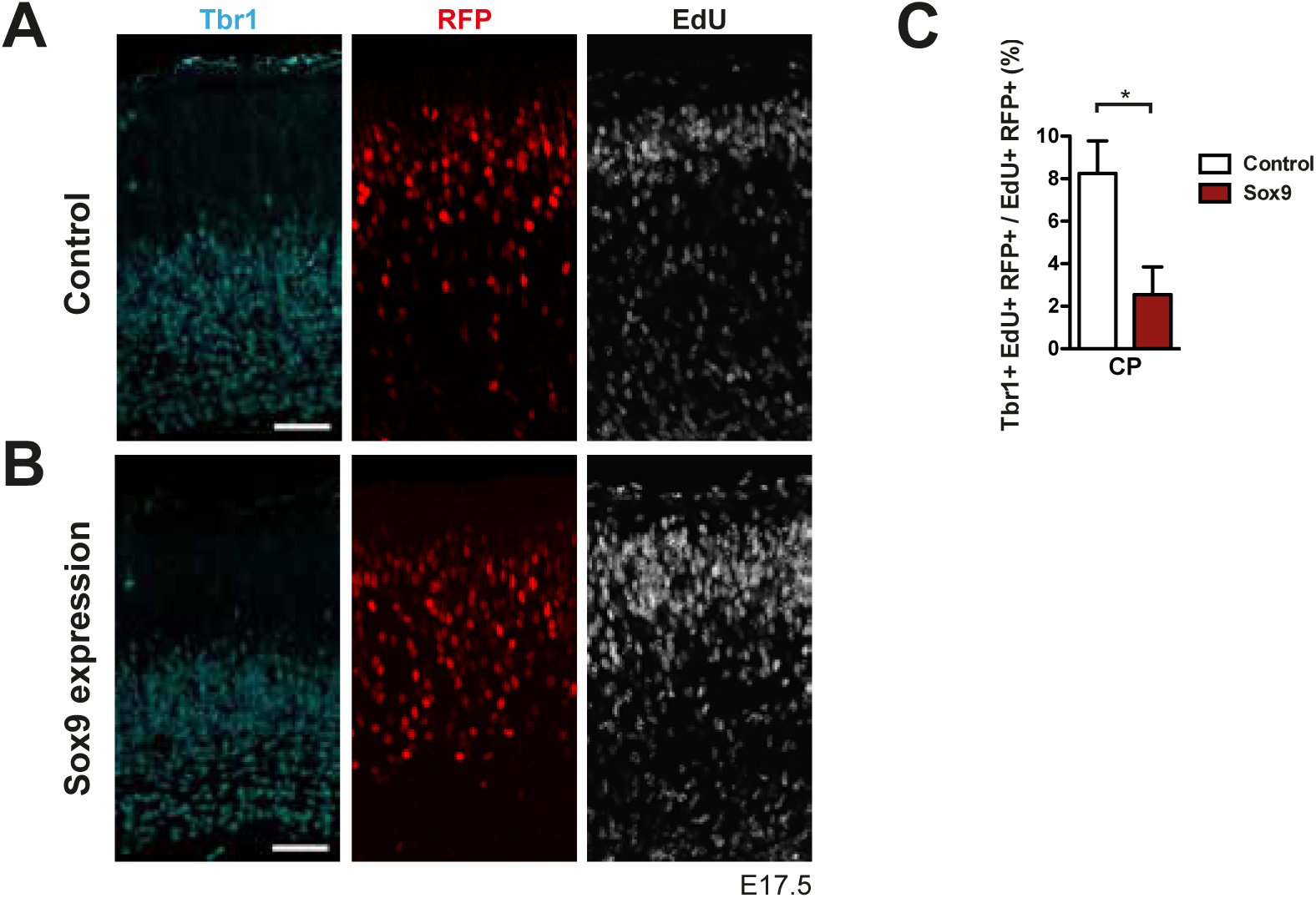
Conditional Sox9 expression results in a decrease of Tbr1-positive neurons in the progeny of the targeted, EdU-labeled mouse BPs. Heterozygous *Tis21*-CreER^T2^ mouse embryos received tamoxifen at E12.5 and E13.5, and the neocortex was *in utero* electroporated at E13.5, followed by triple (immuno)staining for Tbr1 (cyan), RFP (red) and EdU (white) of the CP, 4 days after electroporation of either control construct (**A**) or conditional Sox9 expression construct (**B**) (see Fig. 3A). A single pulse of EdU was administered at E14.5, i.e. 24 hours after electroporation and 3 days prior to analysis. **(C)** Quantification of the percentage of EdU and RFP double-positive nuclei in the CP that are Tbr1-positive, i.e. the percentage of Tbr1-positive neurons in the progeny of the targeted, EdU-labeled mouse BPs, 4 days after electroporation of control construct (white column) or conditional Sox9 expression construct (red column) and 3 days after EdU administration at E14.5. **(A, B)** Scale bars, 50 µm. **(C)** Two-tailed, unpaired *t*-test: * p<0.05. Data are the mean of 3 embryos, each from a different litter; for each embryo, two microscopic fields, each of 200-µm apical width, were counted, and the values obtained were averaged. Error bars represent SEM.

**Supp. Table 1. Lists of genes whose expression is up- or downregulated 48 hours after conditional Sox9 expression.**

**Tab-1 (Sox9_up_PC):** List of genes that are upregulated upon conditional Sox9 expression. **Tab-2 (Sox9_down_PC):** List of genes that are downregulated upon conditional Sox9 expression. Columns in each tab denote the following information consecutively: **A:** Ensemble gene ID, **B:** Gene name, **C:** Description, **D:** Average count for Sox9 sample, **E:** Average count for control sample, **G:** Logarithmic fold change (FC) of counts between Sox9 and control samples, **J:** p-value, K: adjusted p-value, **W:** Average FPKM value for control sample, **X:** Average FPKM value for Sox9 sample, **Y:** Fold change (FC) between average FPKM values of Sox9 and control samples, **Z:** Logarithmic fold change (FC) between average FPKM values of Sox9 and control samples.

**Supp. Table 2. Datasets of RNA-sequencing and gene ontology (GO) term enrichment analysis of Sox9 and control samples.**

**Tab-1 (FPKM):** FPKM values for each replicate of Sox9 and control samples. **Tab-2 (GO_BP):** Gene ontology (GO) term enrichment analysis for biological process (BP). **Tab-3 (GO_CC):** Gene ontology (GO) term enrichment analysis for cellular component. **Tab-4 (KEGG):** KEGG pathway analysis.

